# Structural Basis of Serine Protease Inhibition by Antibodies from Biased Fab Phage-Display Libraries

**DOI:** 10.64898/2026.03.12.711446

**Authors:** Kyle J. Anderson, Melody S. Lee, Natalia Sevillano, Gang Chen, Michael J. Hornsby, Sachdev S. Sidhu, Charles S. Craik

## Abstract

Biased Fab phage-display libraries were designed to determine whether inhibitory CDR H3 motifs from potent anti-matriptase antibodies could be transferred to target homologous serine proteases. Using reverse-binding and substrate-like H3 motifs from parental clones A11 and E2 as templates, six synthetic libraries with 10^10^ diversity were constructed. Selection against matriptase identified sixteen inhibitors with sub-100 nM potency, representing 100,000-fold improvement over circularized H3 loops alone. Selection against TMPRSS2, a serine protease implicated in viral entry and prostate cancer with 43% sequence identity to matriptase, yielded binders with micromolar inhibitory potency. Selection against urokinase plasminogen activator (uPA, 35% identity) identified binders that adopted a substrate-like CDR H3 binding mode in our structural models. Across all reference structures, including the separately identified uPA inhibitor AB2 (PDB: 9PYF, deposited with this work), benchmarking of five co-folding methods and rigid-body docking showed that co-folding consistently achieved acceptable to high quality DockQ scores, outperforming traditional docking and capturing the recognition of key active site determinants. Ensemble predictions of mutational binding energy changes (ΔΔ*G*) using these models identified key paratope-epitope interactions, with predictions validated through mutagenesis. This work establishes a framework integrating biased antibody libraries with computational structure prediction and analysis for targeting conserved protease epitopes.

## 1 Introduction

Developing specific inhibitors against individual proteases remains a central challenge for both tool-compound and therapeutic applications. Structural studies of trypsin-like proteases have extensively characterized the principles governing serine protease activation, substrate recognition, and inhibition by natural protein inhibitors (Huber and Bode, 1978; Bode and Huber, 1992, 2000), revealing that members of a given family share highly conserved active sites. This conservation is particularly evident among type-II transmembrane serine proteases (TTSPs), where alignment of available protease domain structures shows near-identical active site architecture despite more diverse surface loop geometries arising from differences in loop length and sequence (Goettig et al., 2019). The resulting substrate specificities are correspondingly similar, with TTSPs sharing a strong preference for basic residues in the P1 position directly N-terminal to the scissile bond (Rawlings et al., 2016; Barré et al., 2014). Consequently, small molecule and macromolecular inhibitors designed from cognate substrates of matriptase and TMPRSS2 often cross-react with closely related family members (Han et al., 2016; Mahoney et al., 2021; Damalanka et al., 2024; Kojima et al., 2008; Steinmetzer et al., 2006; Stoop and Craik, 2003). Recent substrate profiling and structure–activity relationship campaigns employing unnatural amino acids have achieved potent and selective inhibition of HGFA, matriptase, and hepsin, though this required a significant medicinal chemistry effort (Mahoney et al., 2024). Inhibitory antibodies offer an alternative route to specificity: their large binding surfaces, composed primarily of hypervariable complementarity determining region (CDR) loops, sample vast conformational space and can form contacts spanning hundreds to thousands of square angstroms, enabling both high affinity and selectivity for complex three-dimensional epitopes.

TMPRSS2 and urokinase plasminogen activator (uPA) represent therapeutically important members of the serine protease family that share significant active site homology with matriptase. TMPRSS2 has emerged as a therapeutic target due to its essential role in SARS-CoV-2 spike protein priming and viral entry (Hoffmann et al., 2020; Shapira et al., 2022), as well as its association with aggressive prostate cancers through androgen-regulated expression and proteolytic activation of pro-hepatocyte growth factor (Lin et al., 1999; Lucas et al., 2014). Similarly, uPA is implicated in tumor invasion and metastasis through its activation of plasminogen and degradation of extracellular matrix components, making it a target for anti-metastatic therapies (Andreasen et al., 2000; Salamouni et al., 2022; Su et al., 2016; Masucci et al., 2022). Alignment between protease domains shows matriptase shares 43% sequence identity with TMPRSS2 (60% similarity) and 35% identity with uPA (55% similarity), with particularly high conservation of amino acids in the substrate binding cleft and active site. These relationships suggested that inhibitory motifs effective against matriptase might be transferable to these related proteases.

Previously, we and others identified several antibody-based inhibitors against matriptase, with some of them exhibiting picomolar affinities (Farady et al., 2008, 2007; Foltz et al., 2005; Schneider et al., 2012; Sun et al., 2003; Tamberg et al., 2019), that have been used for molecular imaging and interrogating matriptase activity and overexpression in cancer (Darragh et al., 2010; LeBeau et al., 2013). Crystal structures of the fragment antigen binding (Fab)–protease complex elucidated the mechanism of matriptase (MT-SP1) inhibition by clones A11 and E2 where the antibody CDRs imparted inhibitory specificity and potency by occluding the active site. The main drivers of inhibition were the long CDR heavy chain 3 (H3) loops, containing key arginine residues that were inserted into the protease active site in either a native “substrate-like” or “reverse-binding” orientation, where cleavage of the antibody loops was prevented: **Figure 1A** (Farady et al., 2008). The picomolar affinities achieved by A11 and E2 represent exceptionally potent antibody–protease interactions, likely arising from extensive contacts between multiple CDR loops and protease surface features beyond the active site (Schneider et al., 2012). This suggests that while the CDR H3 motif provides the core inhibitory interaction through active site occlusion, flanking CDR contacts with protease surface loops contribute substantially to overall binding affinity and specificity over other protease family members (Ganesan et al., 2010).

**Figure 1:**
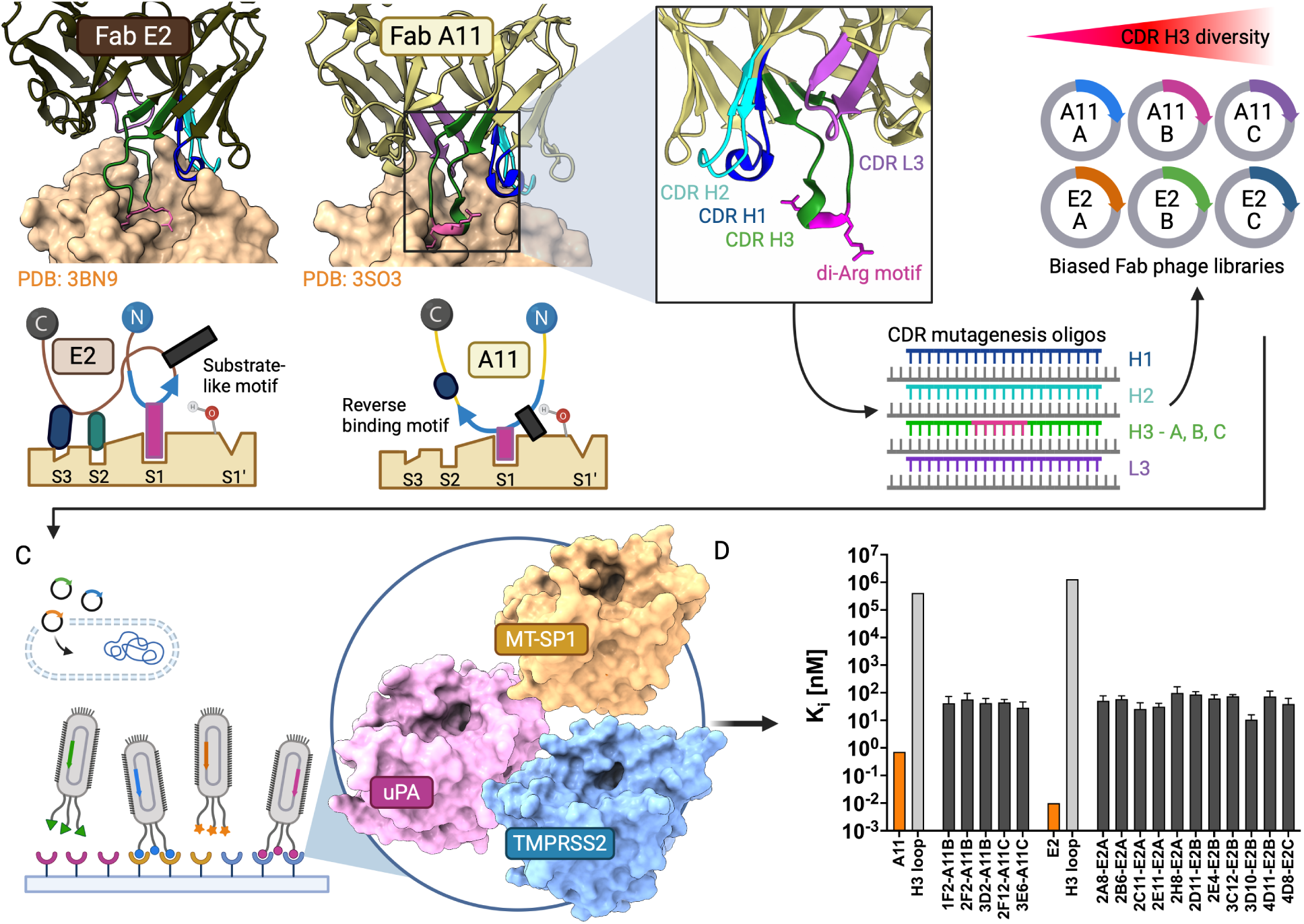
Biased library construction and validation against matriptase. (A) Crystal structures of inhibitory CDR H3 motifs from parental anti-matriptase clones A11 and E2 with cartoon models highlighting factors that prevent cleavage: the reverse orientation of the H3 loop of A11, E2 with kinked H3 loop positions backbone away from catalytic serine residue with hydroxyl shown. Also shown are matriptase substrate pockets S3, S2, S1, and S1*^′^*. (B) CDRs H1 (blue), H2 (teal), H3 (forest green), and L3 (purple) where library was diversified by separate oligo pools (oligos in **Supplement 1**); minimal di-basic arginine motif (magenta). (C) Phage display selection of biased libraries against matriptase, TMPRSS2 and uPA. (D) Inhibition of the 16 biased library hits against matriptase with low nM *K*_I_s (dark grey) vs. parental A11, E2 (orange) and circularized H3 loops alone (light grey). Created in BioRender.

There are several other inhibitory antibodies that target serine proteases at varying stages of development which employ a variety of mechanisms to achieve inhibitory potency (Farady and Craik, 2010). These mechanisms include the use of other CDR loops besides H3 for active site targeting, though many employ non-active-site-directed or allosteric mechanisms. Notable examples include an allosteric anti-tryptase antibody for severe asthma treatment (Maun et al., 2019), allosteric bi-specific targeting of kallikrein 5 and kallikrein 7 (Chavarria-Smith et al., 2022), active-site-directed targeting of plasma kallikrein and kallikrein 2 (Kenniston et al., 2014; Pandit-Taskar et al., 2024), a non-active-site-directed uPA inhibitor that triggers rezymogenation (Jiang et al., 2012), and an anti-XIa antibody that uses CDR L1 and L3 in an active-site-directed manner (Ely et al., 2018). The effective use of other CDR loops such as L1 is of particular interest to this study. Recently, the structure of AB2, an affinity matured clone from the competitive uPA inhibitory antibody U33, was solved in complex with uPA. These anti-uPA clones had much shorter CDR H3 loops compared to the matriptase inhibitors; X-ray footprinting mass spectrometry (XFMS) and X-ray crystallography showed that these anti-uPA antibodies used extended CDR L1 loops as the basis of their inhibitory potency (Sevillano et al., 2021).

In this work, we determined if biasing phage library diversity with inhibitory CDR H3 loop motifs from our previously identified anti-matriptase Fabs could aid in the identification of antibody-based binders with potential inhibitory activity against other serine proteases. Our approach was motivated by the established correlations between the sequence composition of antibody CDRs and the structural nature of the antigen. Such relationships underpin the successes of prior attempts at ‘biasing’ antibody library diversity to recognize specific structural features (Collis et al., 2003; Persson et al., 2006; Almagro, 2004). Recently these works have progressed from the first libraries ‘focused’ to recognize convex sites (Persson et al., 2006), to those now targeting high-value antigen families such as GPCRs (Liu et al., 2021a). We believe using structurally informed diversity to target the serine protease family active-site will improve the hit-rate and further enhance the discovery power of well-established *in vitro* selection methods (Fellouse et al., 2004, 2007; Fellouse and Sidhu, 2005; Sidhu and Fellouse, 2006).

Our primary aim was to establish whether the CDR H3 inhibitory motif could be successfully repurposed to target homologous proteases, not to achieve affinities comparable to the exceptional parental clones A11 and E2. The diversification of CDRs L3, H1, and H2 required for library construction was expected to disrupt some of the peripheral interactions responsible for the picomolar parental potency. We defined success as identifying binders that retained the H3-directed binding mode, even at substantially reduced affinity, thereby validating the biased library concept for protease family targeting. Subsequent affinity maturation would require separate optimization campaigns tailored to each target protease.

In addition, given that experimental structure determination was not feasible for our biased library hits due to their modest affinities, we also sought to determine whether computational structure prediction methods could provide sufficient accuracy to rationalize our experimental observations. Several protein co-folding methods are now available, including AlphaFold2-multimer (Evans et al., 2022; Jumper et al., 2021; Mirdita et al., 2022), AlphaFold3 (Abramson et al., 2024), Boltz-2 (Passaro et al., 2025), Chai-1 (Discovery et al., 2024), and OpenFold3 (The OpenFold3 Team, 2025), alongside established docking tools such as ClusPro (Kozakov et al., 2017). These architecturally distinct approaches provided an opportunity to benchmark prediction accuracy across methods and assess whether accurate modeling reflects structural features of this epitope family rather than method-specific artifacts. We further examined whether computational ΔΔ*G* predictions using these modeled structures could retrospectively explain and contextualize the outcomes of our *in vitro* mutagenesis campaigns and inform future library design. Recent advancements in ΔΔ*G* prediction methods have leveraged ML models to improve both accuracy and speed over traditional physics-based methods like Rosetta (Barlow et al., 2018) and FoldX (Delgado et al., 2019). Building upon these recent findings, we present an ensemble workflow to run and aggregate predictions from physics-based methods alongside new ML models including mCSM-AB2 (Myung et al., 2020), GeoPPI (Liu et al., 2021b), and GearBind (Cai et al., 2024). Ultimately, we found that co-folding methods, benchmarked against crystal structures and a rigid-body docking baseline, alongside the results from our *in vitro* selections, provided a structural framework for understanding the binding modes of antibodies from our biased libraries and retrospectively identifying factors that could limit affinity maturation success.

## 2 Results

### 2.1 Biased library design and validation against matriptase

To examine the effectiveness of the inhibitory CDR H3 loops from anti-matriptase antibodies for more general protease inhibition, we constructed new phage display libraries, termed “biased libraries”. These libraries were based on the previously characterized reverse-binding and substrate-binding motifs from clones A11 and E2, **Figure 1A**. Designed degenerate oligonucleotides were used to introduce diversity at four of the six CDRs that were previously identified as the most important for antibody–antigen interactions in the parental anti-matriptase binders A11 and E2 (Oligos in **Supplement 1**). In this library, CDR light chain 1 (L1) and light chain 2 (L2) were unchanged from the template. Diversity mimicking the amino acids found in natural binding sites was introduced at CDR L3, while limited diversity was introduced at CDR H1 and H2 loops (**Figure 1B**). For the added diversity around the critical reverse-binding and substrate-like motifs on CDR H3, three sub-libraries were built from the H3 loops of A11 and E2 with varying degrees of wild type conservation. Sub-library A contained the most conserved sequence from the parental motif, sub-library B contained a conserved six to seven residue turn motif, and sub-library C preserved only the minimal di-arginine motif. Since the basic arginine residue at position 100b is important for serine protease inhibition, the di-arginine motif was the minimal requirement in the design of CDR H3 in the biased library (Farady et al., 2008). The diversity of each sub-library was determined by titering to be around 4 × 10^9^ (all antibody sequences numbered in Chothia scheme).

To test the design of the biased library, we performed selections against matriptase and panned with each of the six sub-libraries separately, **Figure 1C**. After three rounds of panning, thirty hits with unique sequences in the CDR H3 loop clones were identified from all the sub-libraries. An enrichment of hits from the B sub-library was observed for both A11 and E2. All antibodies were expressed and purified for further characterization. IC_50_ and relative *K*_I_ values were calculated for the clones identified, and 16 out of 30 (53%) showed double-digit nanomolar inhibition of matriptase activity (**Figure 1D**) (CDRs in **Supplement 1**). The heavy and light chains for each antibody were sequenced and longer loop lengths (1–2 residues) in CDR H3 were observed (14/16 clones).

Although each sub-library was panned independently, the resulting hits showed mutations in multiple CDRs because diversity was introduced at CDR L3, H1, and H2 in addition to H3 during library construction (**Supplement 1**). The sub-library designation (A, B, C) refers only to the degree of H3 conservation, not to the other diversified CDRs. The observation that longer H3 loops were enriched during selection (14/16 hits) may reflect the different surface loop topologies between matriptase and the parental A11/E2 interaction surfaces, or may indicate that additional loop length provides conformational sampling advantages during selection. Loop length diversity of ±1–2 residues was included for CDR H3 in the initial library design but not for other CDRs. The most potent antibodies containing the substrate-binding motif came from the more conserved E2 sub-libraries. By contrast, the most potent antibodies with the reverse-binding motif came from the less conserved A11 sub-libraries.

These results are consistent with the relative contributions of heavy and light chain buried surface area in the parental antibody–matriptase complexes, where E2 derives more binding energy from its conserved H3 motif while A11 relies more heavily on peripheral CDR contacts. The inhibition values of these biased library hits are approximately 100-fold less potent than the original parental A11 and E2, consistent with our expectation that CDR diversification would disrupt some peripheral interactions responsible for parental potency. Importantly, these biased library hits retain 100,000-fold improved potency over the circularized H3 loops alone (**Figure 1D**), demonstrating that the antibody scaffold substantially enhances the inhibitory capacity of the H3 motif even when other CDRs are diversified. This validates the core premise that the H3 inhibitory motif can function within a diversified antibody framework (Schneider et al., 2012).

### 2.2 Computational modeling of biased library antibody complexes

We used multiple structure prediction and docking methods to provide a structural framework for the binding of antibodies from our biased library. Drawing from methods used for the Critical Assessment of Predicted Interactions (CAPRI), DockQ is a standard metric for assessing the quality of protein interfaces within modeled complex structures and was used to determine the accuracy of the predicted models against given reference structures (Mirabello and Wallner, 2024). DockQ scores reflect an average of the fraction of native contacts (*f*_nat_), scaled interface RMSD (iRMSD), and scaled ligand RMSD (LRMSD) and have historically been used to classify models according to the CAPRI metrics (“Incorrect” for scores less than 0.23, “Acceptable” for scores between 0.23 and 0.49, “Medium” for scores between 0.49 and 0.8, and “High Quality” for scores above 0.8 (Collins et al., 2024)). Models in the acceptable accuracy tranche can include moderate deviations from the reference structure including up to 10 Å of LRMSD, where medium and high accuracy cutoffs reflect more stringent cutoffs such as iRMSD *<* 2.5 Å and represent near native models (Chen et al., 2003).

We benchmarked five co-folding methods (AlphaFold3 (AF3) (Abramson et al., 2024), Boltz-2 (Passaro et al., 2025), Chai-1 (Discovery et al., 2024), AF2-multimer (Jumper et al., 2021; Mirdita et al., 2022), and OpenFold3 (The OpenFold3 Team, 2025)) alongside ClusPro rigid-body docking (Kozakov et al., 2017) against crystal structures of A11–matriptase (PDB: 3SO3) and E2–matriptase (PDB: 3BN9). For AF3, Boltz-2, and Chai-1, we ran 100 predictions per complex with unique seeds; for AF2-multimer and OpenFold3, we ran five predictions per complex with unique seeds (templates disabled for all methods). We report the top-ranked model by each method’s confidence metric. We acknowledge that the crystal structures of A11 and E2 were present in training datasets for several methods; to avoid trivial template-based retrieval, we disabled template usage for all predictions and relied solely on each model’s learned weights. Separate runs with templates enabled showed improved scores for AF2-multimer, most dramatically for E2–matriptase (DockQ 0.47 to 0.94), while AF3 showed negligible template benefit (**Supplement 3**), consistent with differences in how each architecture leverages structural information. For the matriptase complexes, Boltz-2 achieved the highest accuracy (DockQ 0.91 and 0.89 for E2 and A11, both High Quality). AF2-multimer scored 0.47–0.72 (Acceptable to Medium), AF3 scored 0.37–0.60 (Acceptable to Medium), Chai-1 scored 0.47–0.58 (Acceptable to Medium), and OpenFold3 scored 0.44–0.64 (Acceptable to Medium). ClusPro rigid-body docking produced Acceptable-quality models (0.36–0.37), with all co-folding methods outperforming the docking baseline, **Figure 2B**. AF3 and Chai-1 showed larger gaps between rank-1 and best-sampled scores (up to 0.10 and 0.19 DockQ respectively), suggesting that increased sampling can improve model selection for these methods (**Supplement 3, continued**).

**Figure 2:**
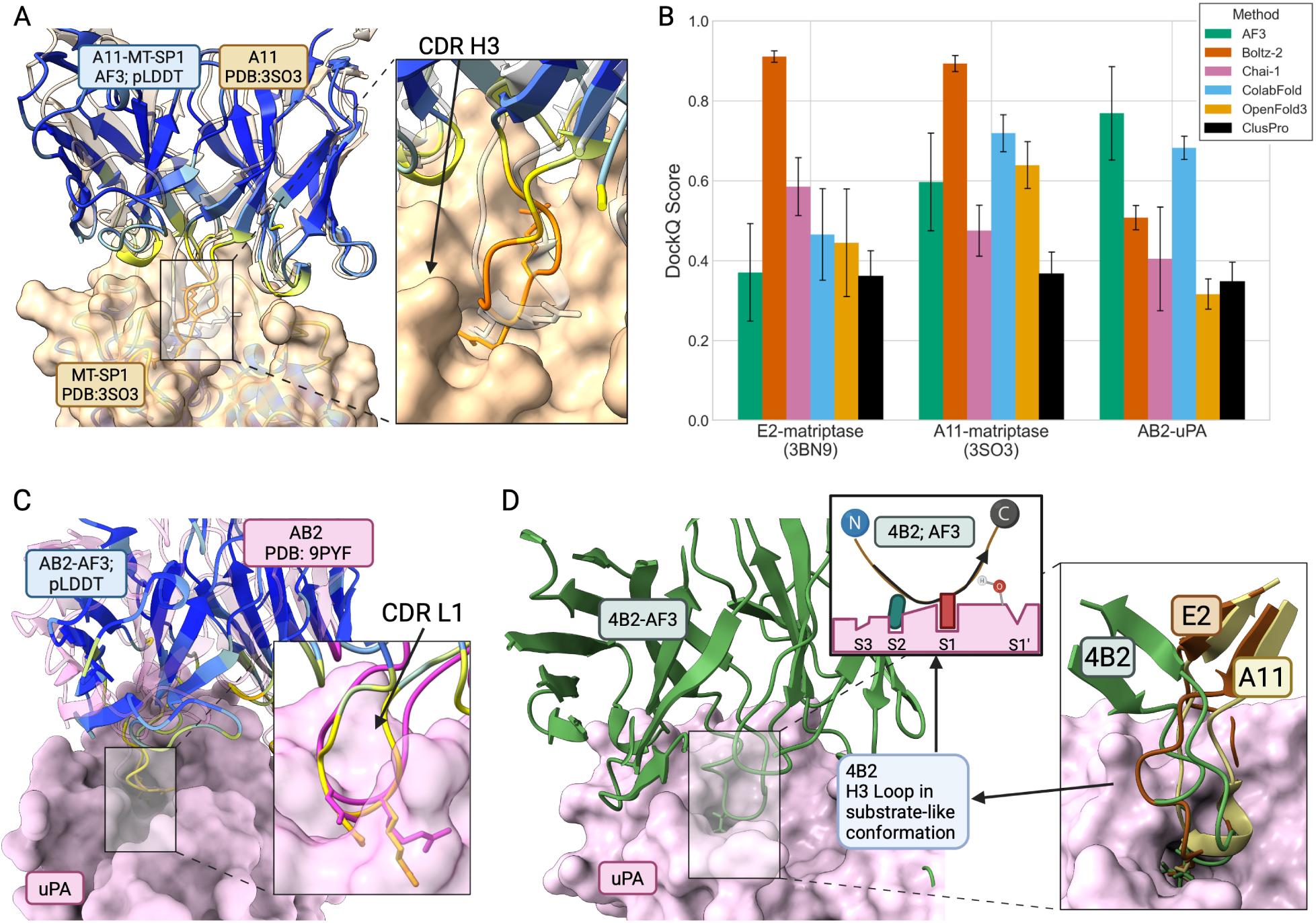
Computational models capture CDR H3 and L1 directed inhibitory modes. (A) AF3 model of parental anti-matriptase clone A11, colored by pLDDT with crystal structure of A11 in complex with MT-SP1 (transparent grey), PDB: 3SO3. (B) Comparison of mean DockQ scores across six structure prediction and docking methods (no templates) for antibodies with known structures. Error bars: s.d. across prediction seeds or docking clusters. (C) AF3 model of anti-uPA inhibitor AB2 (colored by pLDDT) aligned to AB2–uPA crystal structure (transparent pink). (D) biAb-uPA-1, modeled in complex with uPA using AF3; CDR H3 in substrate-like conformation in protease active site. All pLDDT coloring schemes in the standard DeepMind format: very high confidence (dark blue), high confidence (light blue), medium confidence (yellow), low confidence (orange). Created in BioRender.

Despite some inaccuracies intrinsic to the problem of modeling antibodies (i.e. CDR loop conformations), all co-folding models accurately localized the CDR H3 loop into the substrate binding cleft and coordinated basic CDR H3 residues within the protease S1 site, indicating that these inhibitors from the biased libraries retain the H3 directed inhibitory basis from the parental Fabs A11 and E2. Examining confidence metrics for these models, predicted Local Distance Difference Test (pLDDT) shows most antibody framework regions are modeled with very high confidence (pLDDT *>*90) with most CDR loops falling into ‘good confidence’ (90 *>* pLDDT *>* 70), and and CDR H3 the lowest with pLDDT ∼60 in some regions, **Figure 2A**. The Predicted Aligned Error (PAE) domains separate the antibody from the protease and parts of the CDR H3 loop, indicating good intra-domain confidence but more uncertainty in their relative orientations. Predicted template modeling (pTM) and interface predicted template modeling (ipTM) scores were all above 0.8 except for E2 models which scored lower, ∼0.6 (additional AF metrics in **Supplement 3**). Overall, these results indicate that predictions of the separate antibody and protease conformations are relatively accurate, with more uncertainty in the precise antigen-binding pose.

With the biased library validated against matriptase for identifying nanomolar inhibitors, we sought to examine how our library might be used to identify novel inhibitors for other closely related proteases, namely TMPRSS2 and urokinase (uPA). Both TMPRSS2 and uPA share the same chymotrypsin fold as matriptase and have homologous active site pockets; alignment between protease domains of human uPA and MT-SP1 shows 35% sequence identity and 55% similarity, based on the BLOSUM62 matrix, TMPRSS2 is more closely related with 43% identity, 60% similarity.

Our recently deposited crystal structure of the previously reported inhibitory antibody AB2 in complex with uPA (PDB: 9PYF) provided additional validation for the accuracy of structure prediction on this epitope family. With the separate inhibitory mode of AB2 utilizing CDR L1, we found that the accuracy of co-folding methods on the serine protease active site epitope extended beyond just the CDR H3-biased interactions from the original matriptase inhibitors, to include uPA inhibitors that instead employ an extended CDR L1 loop which coordinates an Asp residue in the protease active site, **Figure 2C**. AB2’s use of CDR L1 as the basis for serine protease inhibition offers an additional explanation for the lack of binders with inhibitory potency against uPA discovered from our CDR H3-biased libraries. For the AB2–uPA complex, AF3 achieved the highest score (DockQ 0.77, Medium), followed by AF2-multimer (0.68, Medium), Boltz-2 (0.51, Medium), Chai-1 (0.40, Acceptable), ClusPro (0.35, Acceptable), and OpenFold3 (0.32, Acceptable), **Figure 2B**. The AB2–uPA structure was deposited after the training data cutoffs for all methods, providing validation that accurate modeling of this epitope family generalizes to structures not seen during training. The consistency of binding-mode prediction across six independent methods suggests that the serine protease active site represents a tractable structural target for current co-folding methods. The performance of these methods in predicting the binding mode of AB2 provided confidence that antibody models with this epitope family can be accurate enough to structurally inform the interpretation of selection outcomes.

### 2.3 Selection of the biased library against uPA

Selection of our biased library against active, recombinant biotinylated uPA (LWMuPA-C122A) was performed using streptavidin magnetic beads for three rounds of selection. Five unique clones that bound uPA were identified from our biased libraries. Four of these hits showed binding to uPA in the high nanomolar range as measured by biolayer interferometry (BLI), however, none had any inhibitory effect against uPA at a concentration of 1 *µ*M. Three of these biased library hits even behaved as substrates for uPA, with cleavage confirmed by SDS-PAGE after incubation; see supplemental information in (Sevillano et al., 2021).

We then used AF models to provide a structural basis for these results against uPA. Models of biAb-uPA-1, one of the biased library hits where cleavage was observed by SDS-PAGE, showed a binding mode where an extended CDR H3 was presented to the enzyme active site pocket. This interaction was similar to the characteristic active site insertion of the CDR H3 motif observed in the parental anti-matriptase clones A11 and E2. Particularly, biAb-uPA-1 showed its H3 loop interacting in a substrate-like orientation with the uPA active site, unlike E2, whose substrate-like presentation of this loop avoids cleavage by adopting a unique kinked conformation that positioned the backbone away from the catalytic serine. biAb-uPA-1 likely lacked this unique positioning of CDR H3 resulting in the observed cleavage. These AF models ultimately provided a structural rationale for the lack of inhibitory potency and cleavage of the biased library hits, such as biAb-uPA-1, against uPA, **Figure 2D**.

The observation that our H3-biased libraries yielded binders that potently recognized the uPA active site, yet adopted a substrate-like binding mode resulting in cleavage rather than inhibition, indicated that the CDR H3 motif alone was insufficient for achieving uPA inhibition. This finding motivated continued efforts to identify inhibitory antibodies against uPA, ultimately leading to the identification and characterization of AB2, which achieves inhibition through an extended CDR L1 loop that resists proteolytic cleavage (Sevillano et al., 2021). Together, these outcomes underscore that different proteases may be optimally targeted by different CDR loop architectures, and that the unique surface topology of uPA around the active site may favor L1-directed approaches over H3 insertion.

### 2.4 Selection of the biased library against TMPRSS2

Active, recombinant TMPRSS2 enzyme was biotinylated and attached onto streptavidin-coated plates or beads for the selection with the biased libraries. Three rounds of selection were sufficient to observe enrichment of hits against TMPRSS2, and seventeen unique sequences in the CDR H3 loop were identified. These hits came mainly from the least conserved sub-library C for both inhibitory motifs. Four clones (biAb-TM-1, biAb-TM-7, biAb-TM-8, and biAb-TM-5) were expressed and purified for further characterization. biAb-TM-5 and biAb-TM-8 showed moderate inhibition of TMPRSS2 activity with IC_50_ values in the *µ*M range. biAb-TM-7 was tested in an IgG format and achieved an IC_50_ of 640 nM (**Supplement 2**).

When examining the AlphaFold models for the biased library hits against TMPRSS2 such as biAb-TM-5, the predicted binding poses all retained the critical CDR H3 mediated interactions with the protease substrate binding pocket, coordinating an arginine residue with the conserved aspartic acid at the base of the S1 site. Compared to models of the matriptase inhibitors, these models show the biased anti-TMPRSS2 clones approaching the protease from a slightly different angle and orientation, likely due to differences in the protease surface topologies such as the shorter 60s surface loop of TMPRSS2 (**Figure 3B, C**). These models further suggest that the antibodies derived from our biased libraries retain the CDR H3-biased interactions, and that the original inhibitory interactions can be effectively translated to a homologous protease.

**Figure 3:**
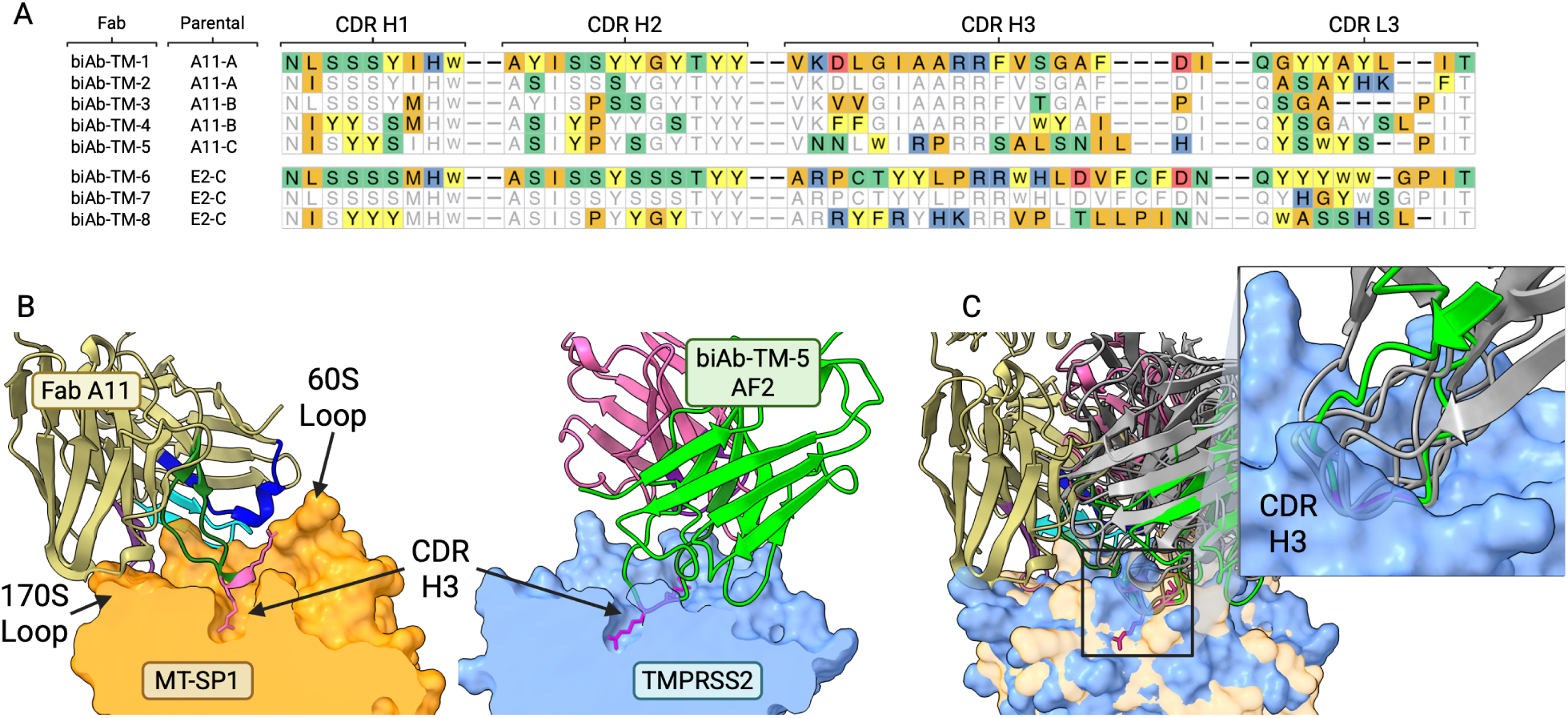
TMPRSS2 inhibitors from biased library. (A) Alignment of CDRs for Biased Library TMPRSS2 hits; grey positions retain parental residues; mutant positions are colored by residue type. (B) Cut-away of A11–matriptase crystal structure and biAb-TM-5–TMPRSS2 AF2 model. (C) Overlay between A11 crystal structure with matriptase and AF2 models of biased library TMPRSS2 hits with H3-directed binding modes (inset) (biAb-TM-5 in green and pink; other biAb-TM-X models in grey). Created in BioRender.

The observation that TMPRSS2 binders were identified primarily from the least conserved sub-library C, combined with their reduced potency, supports our model that diversity added to our biased libraries disrupted the specific interactions originally responsible for the exceptionally high affinity and specificity of A11 and E2 for matriptase. However, this potential for new CDR contacts is what can accommodate binding to other serine proteases such as TMPRSS2. The biased libraries thus appear to trade high affinity to a specific protease family member for the ability to generate weaker binders across multiple family members. Based on our observed performance of AF models accurately capturing antibody interactions with uPA and matriptase, we believed these models of the weak TMPRSS2 biased library hits such as biAb-TM-5 could be useful in understanding the structural basis of their binding modes and informing affinity maturation strategies.

### 2.5 Structure-guided mutagenesis of TMPRSS2 inhibitors

To identify areas where the biased library hits could be improved, as well as where the library design can be improved, we initially constructed a rigid-body homology model of biAb-TM-5 in complex with TMPRSS2 using RosettaAntibody (Sircar et al., 2009; Weitzner et al., 2017, 2014), and I-TASSER (Yang and Zhang, 2015), aligned based on the structure of the parental template A11. The rigid-body structural model of the Fab–TMPRSS2 complex (**Figure 4A**) highlights regions in the CDRs where the binding affinity and the specificity could be improved through mutation. The 170s loop on the protease surface can make potential interactions with the CDR L1 loop, however in the original biased library design, L1 was unchanged from the template. We hypothesized that introducing diversity and length variation in this CDR may improve interactions with TMPRSS2. Additionally, among TTSP family members, the 60s loop of the proteases is very diverse in sequence and length and provides an opportunity for achieving specificity (Goettig et al., 2019). Matriptase has four more amino acids in the 60s loop than TMPRSS2 (**Figure 3B**, **Figure 4B**). With this difference in protease loop sizes, our rigid-body model suggested that incorporating additional length variation in the CDR H1 and L1 loops, increasing their length by 1–2 amino acids, may improve interactions with TMPRSS2 and aid in specificity and affinity.

**Figure 4:**
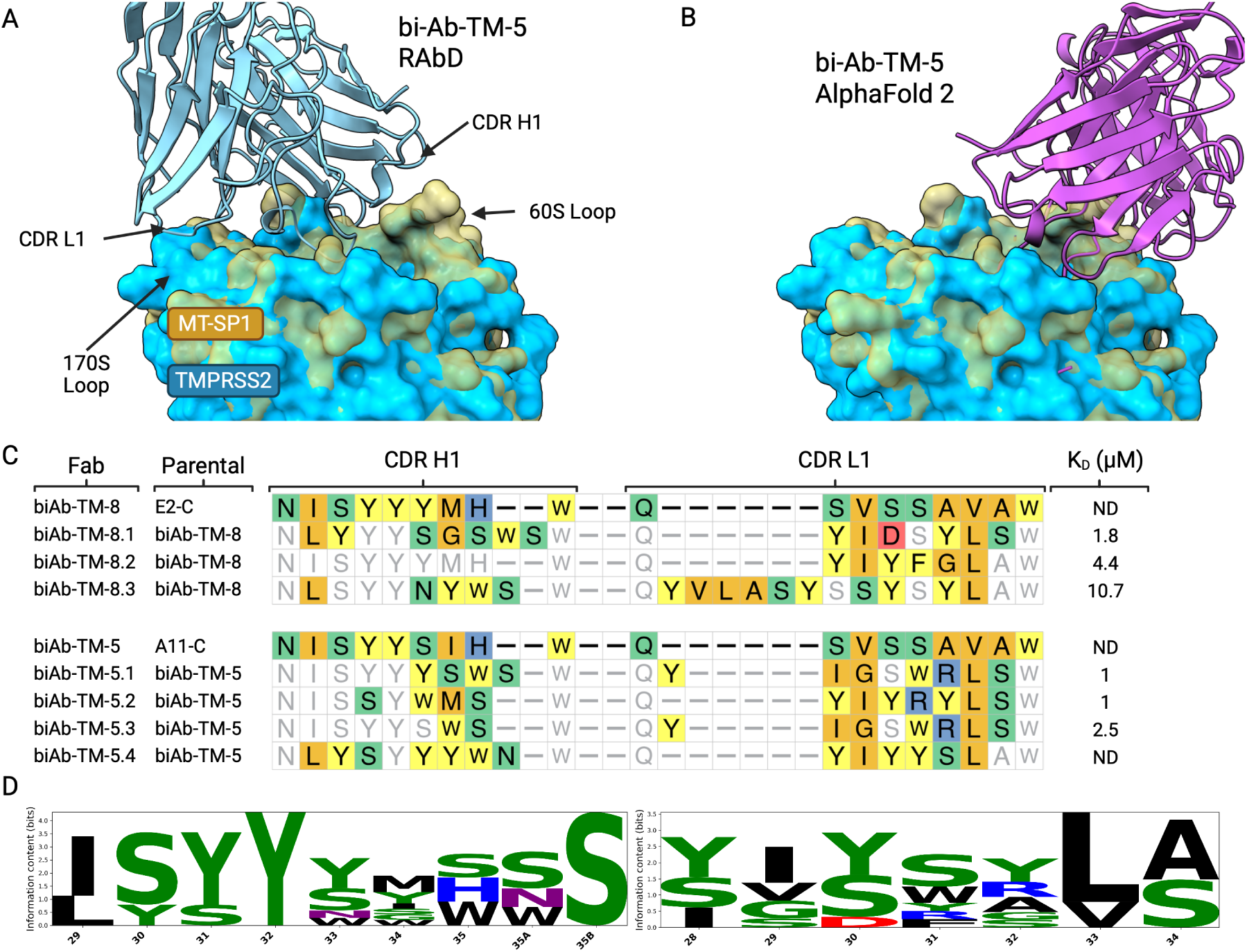
Affinity maturation of biAb-TM-5. (A) Rigid body model from Rosetta Antibody of biAb-TM-5 (light blue ribbons) in complex with TMPRSS2 from I-TASSER; aligned according to PDB 3SO3 A11 model. (B) AF3 model of biAb-TM-5 (magenta) in complex with TMPRSS2. Both protease models colored as blue for TMPRSS2, yellow for matriptase. (C) Alignment of CDRs H1, L1, from affinity matured TMPRSS2 hits with *K*_D_ values from BLI; positions with mutations over the parental clone are colored according to residue chemistry. Residue preference logos for CDR contacts. ND = not determined. (D) Sequence logos for CDRs recovered during biopanning. Created in BioRender.

Using the rigid-body model of the biAb-TM-5 in complex with TMPRSS2 as a guide for improving the biased library, we designed additional degenerate oligonucleotides that introduce length variation and diversity to the CDR H1 and L1 loops (oligos in **Supplement 2**). Clones biAb-TM-1, biAb-TM-7, biAb-TM-8, and biAb-TM-5 were used as the templates onto which diversity was introduced using an optimized site-directed mutagenesis protocol (Chen and Sidhu, 2014), generating a new library with approximately 10^10^ diversity. Two additional rounds of panning against TMPRSS2 using reduced protein concentration and more stringent, longer washes were completed to select for better binders. From the last round, seven hits were selected for additional studies. The seven clones came from two parental templates, biAb-TM-8 and biAb-TM-5. Many of these second-generation hits for TMPRSS2 had longer CDR H1 and L1 loops, an abundance of larger amino acids in the CDR L1 loop (all Fabs), and several contained an additional basic residue in the CDR L1 loop (biAb-TM-5.1, biAb-TM-5.2, and biAb-TM-5.3) (**Figure 4C, D**). Binding affinities for these Fabs, as determined by BLI, revealed *K*_D_ values in the low *µ*M range (**Figure 4C**) and inhibition assays (in IgG format) confirmed their inhibitory potency against TMPRSS2 with IC_50_s in the high nM regime (**Supplement 2**).

### 2.6 ΔΔ*G* predictions

Having completed our H1+L1 mutagenesis campaign using the rigid-body model, we sought to retrospectively examine whether ΔΔ*G* predictions using higher-accuracy AF2 models could explain our selection outcomes and provide guidance for any future library designs. Using the AF2 model of Fab biAb-TM-5 in complex with TMPRSS2 as an input, saturation ΔΔ*G* predictions were performed for all CDR residues using both physics-based methods: FoldX (Delgado et al., 2019) and Rosetta Flex ΔΔ*G* (Barlow et al., 2018), as well ML-based methods: mCSM-AB2 (Myung et al., 2020), GeoPPI (Liu et al., 2021b), and GearBind (Cai et al., 2024). We constructed an ensemble of ΔΔ*G* scores from this array of methods to incorporate the advantages from both physics-based and ML approaches. This combined set of ΔΔ*G* values, with averaged and normalized scores across the CDR positions, is visualized in a heatmap of this energy landscape, **Figure 5A**. Some overall trends in the ΔΔ*G* landscape of biAb-TM-5 bound to TMPRSS2 are readily apparent, such as the dibasic arginine motif on the heavy chain (101a, 101b) where most mutations are scored as highly destabilizing.

**Figure 5:**
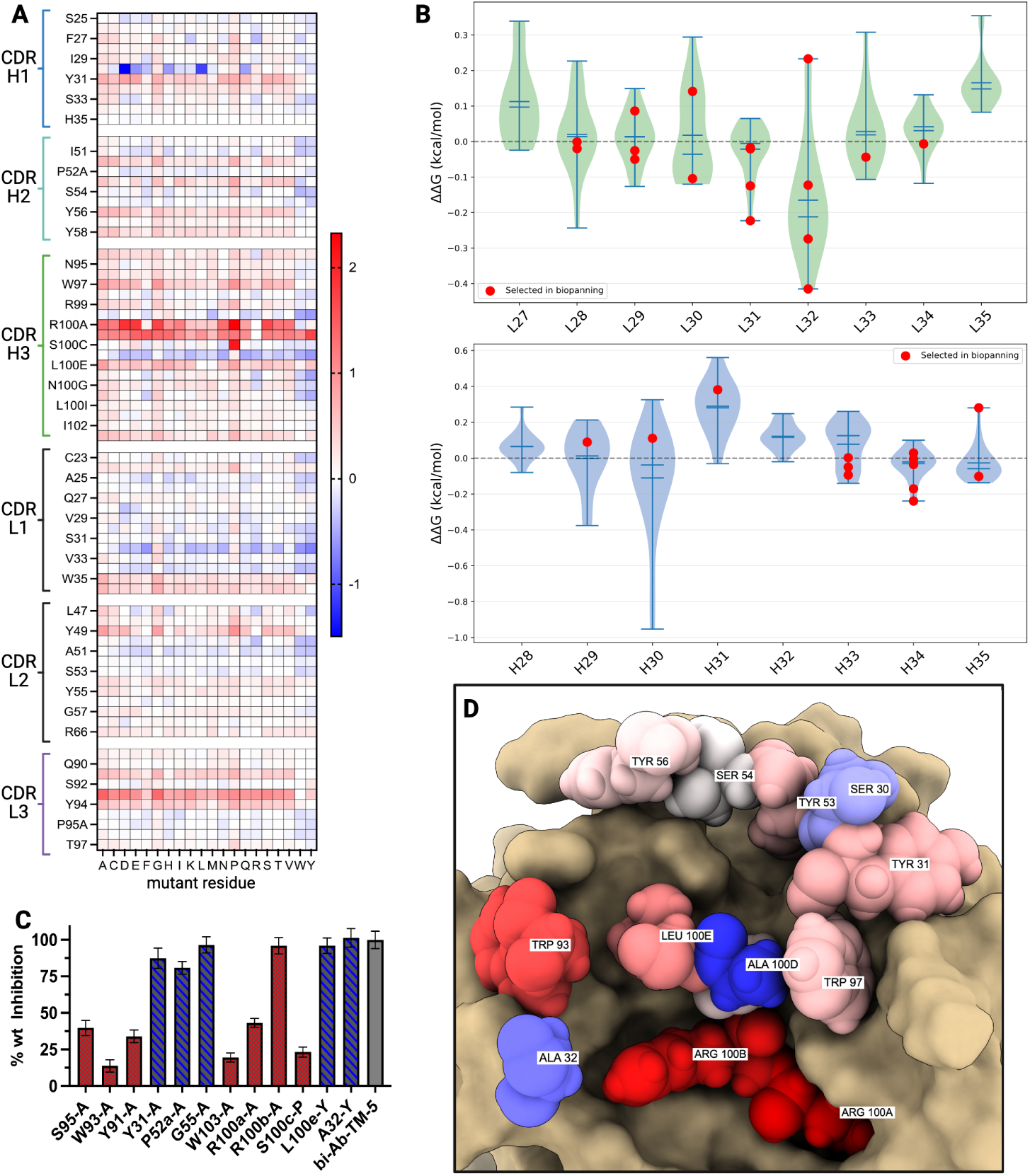
Mutational binding energy landscape of biAb-TM-5–TMPRSS2 complex. (A) PSSM matrix of ΔΔ*G* mutations to CDRs biAb-TM-5; showing average normalized ΔΔ*G* scores (color scheme red – destabilizing, blue – stabilizing). Positions selected for mutagenesis are highlighted with star. (B) Violin plots showing full landscape of ΔΔ*G* values for H1 and L1, negative values indicate stabilizing mutations, red marks indicate ΔΔ*G* of residue mutants identified through biopanning. (C) Percent activity of TMPRSS2 treated with 5 *µ*M of biAb-TM-5 point mutants; percent activity in comparison to the parental biAb-TM-5. Error bars represent standard deviation from *n* = 3 technical replicates. (D) AF2 model of biAb-TM-5 in complex with TMPRSS2 (tan surface) showing CDR residues proximal to protease surface colored by mean ΔΔ*G*. Created in BioRender.

In contrast to our initial rigid-body homology modeling of the biAb-TM-5–TMPRSS2 complex that guided the selection of H1 and L1 as optimal sites for affinity maturation, the ΔΔ*G* scores for H1 suggest that many mutations to this loop are deleterious. In **Figure 5B**, the distribution of ΔΔ*G* scores for both mutated CDRs highlights L1 as having more positions with stabilizing potential; the heavy chain outlier at position 30 was unable to leverage its favorable ΔΔ*G* distribution (mostly for mutations to charged amino acids) as this residue was only diversified in our libraries between tyrosine and serine. Based on these scores, L1 appeared to offer more opportunities for increasing affinity, along with the substitution to bulkier residues in positions flanking the dibasic arginine motif on CDR H3. Per-method ΔΔ*G* heatmaps and position coverage are shown in **Supplement 4**; the average ΔΔ*G* for all CDR positions, along with additional cross-method comparisons are shown in **Supplement 5**.

After selection of our H1+L1 libraries against TMPRSS2, we examined whether mutations identified through *in vitro* biopanning corresponded with our ΔΔ*G* landscape constructed from the AF2 modeled structure. With the red highlights in Figure 5B indicating ΔΔ*G* values of residues selected during this biopanning campaign, qualitative agreement with selection outcomes is apparent. However, formal correlation analysis (Spearman *ρ* = 0.15, *p* = 0.31, **Supplement 6**) did not reach statistical significance due to the limited number of unique mutant sequences identified *in vitro*.

### 2.7 ΔΔ*G* predictions and biAb-TM-5 point mutants’ inhibition of TMPRSS2

With the ability of these ΔΔ*G* scores to provide a rationale for the results of the CDR H1, L1 mutagenesis biopanning, we aimed to further probe the general trends and accuracy of these predictions by making several alanine point mutations to assess the contribution to inhibitory potency of residues at specific positions. biAb-TM-5 heavy chain residues Y31, P52a, G55, W103, R100a, R100b, and light chain residues Y91, W93, S95 were mutated to alanine and their inhibitory effect against commercial TMPRSS2 was measured at a standardized 5 *µ*M Fab concentration and compared against the activity of the parental biAb-TM-5 clone.

These data highlighted alanine mutations at S95, W93, Y91, W103, and R100a as most disruptive to the inhibitory potency of biAb-TM-5 and by proxy its affinity to TMPRSS2. The primarily destabilizing ΔΔ*G* scores for positions like light chain W93 indicate these residues are likely already making important interactions with the protease that are undesirable to disrupt. This was also the case with Y91 and W95, which can be seen along with W93 in **Figure 5D** making contacts with the protease surface and scoring as destabilizing on average.

Several specific point mutations were also made to biAb-TM-5 based on the ΔΔ*G* predictions that include heavy chain S100c-P, L100e-Y, and light chain A32-Y. The activity of these biAb-TM-5 point mutants against TMPRSS2 corresponded to their binding energy predictions with S100c-P significantly reducing potency and L100e-Y and A32-Y both maintaining similar biAb-TM-5 activity. Although L100e-Y and A32-Y were predicted to be stabilizing mutations, the lack of measured increase in inhibitory potency is not unexpected as improving antibody binding often requires multiple mutations (**Figure 5C**).

## 3 Discussion

Antibody engineering has become increasingly reliant on the dynamic application of many different methods for optimizations, many of which are now being performed computationally. The serine protease family provided an ideal test system to demonstrate the accuracy of new *in silico* structure and binding energy prediction methods alongside the continuing importance of structure-guided *in vitro* selection approaches. With the speed and accessibility of these computational methods, our results suggest they can provide a structural framework for interpreting selection outcomes and retrospectively identifying factors that may have limited affinity maturation success. This approach can ultimately contribute to workflows for identifying novel inhibitory antibodies or more broadly, antibodies targeting members within a conserved protein family.

We report the successful use of two different active-site inhibitory motifs around which we designed large synthetic antibody libraries to examine whether such biased diversity could identify binders to related serine proteases. From these biased libraries, we identified binders to matriptase, TMPRSS2, and uPA; demonstrating that the CDR H3 inhibitory motif can be successfully transferred across homologous active site epitopes. The observation that hits from our biased libraries bound all three target proteases, despite the CDR diversification required for library construction, validates the transferability of the CDR H3 inhibitory motif and the biased library concept. This structural understanding of binding modes was ultimately aided by computational models from multiple co-folding and docking methods, which consistently achieved acceptable to medium quality on this epitope family.

Our new matriptase binders were inhibitory, though much less potent than the parental A11 or E2. These affinity and potency differences support our notion that interactions between protease surface loops and other antibody CDRs, besides the active-site binding H3, are responsible for the pM potency of these parental binders (Schneider et al., 2012). As the TMPRSS2 binders from the biased libraries were relatively weak in binding, these tradeoffs in specificity which arise from the diversification of parental CDRs in our biased libraries, likely enabled the successful translation of this inhibitory H3 motif to other proteases in the family. We conclude the diversity added to our biased libraries disrupted the interactions originally responsible for the high affinity and specificity of A11 and E2 for matriptase. However, the new CDR contacts can accommodate binding to other serine proteases such as TMPRSS2 and uPA.

While the uPA binders from the biased library lacked inhibitory activity and were cleaved by the protease, their identification as binders nonetheless validates the transferability of the H3-directed binding mode to this more distantly related target (35% identity). The separate discovery of AB2, achieving inhibitory activity against uPA with an extended CDR L1 loop, is concordant with our understanding that there are many ways antibodies can inhibit serine proteases, not all of which utilize an H3-directed inhibitory mode (Tamberg et al., 2019; Ganesan et al., 2010). We now have a structural understanding of these additional inhibitory motifs against the active site of this protease family, such as those targeting plasma kallikreins (Kenniston et al., 2014; Pandit-Taskar et al., 2024), factor XIa (Ely et al., 2018), as well as uPA. The unique surface topology of uPA around the active site may favor L1-directed approaches over H3 insertion, and future biased library designs for serine protease targets should consider incorporating diversity in multiple inhibitory CDR motifs, not solely CDR H3, to maximize the probability of identifying functional inhibitors against any given family member.

The accuracy of structure prediction models for A11, E2, and AB2 was notable given prior reports of limited success for antibody–antigen complex prediction; Yin et al. (2022) found success rates below 6% at the acceptable accuracy cutoff for their benchmark set of antibody complexes. This discrepancy may reflect the constrained nature of the active-site-directed epitope, where the serine protease substrate binding cleft provides a well-defined structural target compared to the diverse epitopes in general antibody–antigen benchmarks. The limited number of antibody–antigen structures in the PDB for training, combined with the conformational diversity of hypervariable CDR loops, has contributed to the generally modest performance of these models on antibody–antigen systems. Despite these challenges, all five co-folding methods modeled these complexes to acceptable or better accuracies. All co-folding methods outperformed ClusPro rigid-body docking. The strongest multi-method consensus was observed for the AB2–uPA complex, where four of five co-folding methods achieved medium quality despite this structure being deposited after training data cutoffs for all methods. The variance observed across different random seeds and the consistency across architecturally distinct methods further supports genuine prediction rather than memorization of training data. Template-enabled runs with AF2-multimer showed dramatic improvements for A11 and E2 (**Supplement 3**), while AF3 showed negligible template effects, consistent with differences in how each architecture leverages structural information. The success on AB2–uPA demonstrates that accurate modeling generalizes across different CDR-mediated binding modes within this epitope family, not just the H3-directed interactions present in training data. The accuracy of these models supported our interpretation that the biased library binders retain CDR H3-directed binding modes and provided a structural basis for understanding the cleavage observed with the uPA binders.

Our AF models also enabled computational binding energy predictions, a critical component of emerging *in silico* virtual screening approaches. Although the initial mutagenesis of TMPRSS2 hits was guided by a rigid-body homology model, retrospective ΔΔ*G* analysis using AF2 models provided additional insight into our selection outcomes. The ensemble ΔΔ*G* predictions correctly identified the dibasic arginine motif as critical for binding, indicated that CDR L1 offered more mutational opportunities than H1, and qualitatively aligned with biopanning outcomes. These predictions highlight CDR L1 as offering particularly rich mutational opportunity, an insight that is now informing ongoing optimization efforts. The agreement between predicted destabilizing mutations and experimental loss of inhibitory activity provides independent validation that the AF2 model captures essential features of the biAb-TM-5:TMPRSS2 interaction; the alanine scan results for positions W93, Y91, W103, and R100a, all predicted as destabilizing by the ensemble approach, showed the expected reduction in TMPRSS2 inhibition. These results support our perspective that AF models of sufficient accuracy can enable informative ΔΔ*G* analysis, providing a computational framework for understanding selection outcomes and guiding CDR selection for mutagenesis in future campaigns.

The reduced potency of the biased library hits relative to the parental clones A11 and E2 is consistent with our expectations based on the extensive characterization of these exquisitely potent inhibitors and the established contributions of peripheral CDR contacts to their pico-molar affinities (Farady et al., 2008; Schneider et al., 2012; Ganesan et al., 2010). Diversification of these CDRs was a deliberate design choice to broaden target recognition across the protease family, and the observed affinity reductions reflect this intended tradeoff. The computational methods and ΔΔ*G* framework established here provide a foundation for continued optimization of these inhibitors, including further affinity maturation and characterization of selectivity against related serine proteases.

Future development of biased libraries for serine proteases would benefit from incorporating multiple inhibitory motifs (H3, L1, L3) as alternative templates, inclusion of CDR length diversity from the outset, and prospective computational evaluation of target-specific structural requirements before library construction. The workflows established here, combining biased library selection with co-folding structural modeling and ensemble ΔΔ*G* analysis, provide a framework for such iterative improvements. The demonstrated accuracy and cross-method consistency of structure prediction on this epitope family suggests that virtual screening approaches, including prospective ΔΔ*G*-guided library design and computational affinity maturation, warrant investigation for serine protease antibody engineering.

In conclusion, we have developed an integrated approach for identifying antibodies targeting conserved protease active site epitopes. Our approach leverages structural insights from inhibitory CDR motifs for library construction and computational methods for retrospective interpretation of selection outcomes. This strategy has contributed to the identification and characterization of several binding antibodies for TMPRSS2 with modest inhibitory potency, with the potential to enable functional and structural studies to understand its activity and localization in disease. Focused or biased antibody libraries continue to demonstrate themselves as powerful tools for accelerating antibody discovery. Our results suggest the combination of these antigen-tailored *in vitro* methods with emerging *in silico* tools for structure and binding prediction can provide a framework for understanding and improving future antibody development campaigns. The agreement of results across multiple independent prediction methods supports the reliability of the structural insights drawn from these models, with retrospective analysis informing subsequent rounds of library design and selection.

## 4 Materials and methods

### 4.1 Library construction

We introduced these CDR H3 motifs into a high expressing and clinically validated antibody framework VH3-23, VK1-19 that has high stability and reported use for library design (Chen and Sidhu, 2014; Hoet et al., 2005). As the template onto which the library was built, we chose an anti-maltose binding protein framework that was previously used to construct a highly functional naïve synthetic antibody library (Library F) (Persson et al., 2006).

The library was constructed using a phagemid (pHP153) designed for the phage display of a highly stable human Fab framework, as described previously (Chen and Sidhu, 2014). A set of mutagenic oligonucleotides were designed to introduce diversity at CDR L3, CDR H1 and CDR H2 loops (**Supplement 1**). CDR L3 was mutagenized with the oligonucleotides L3-3, 3-4, L3-5, L3-6 and L3-7. CDR H1 was mutagenized with oligonucleotide H1a-4 and CDR H2 was mutagenized with H2a. The template Fab CDR H3 was replaced with the CDR H3 sequence of A11 or E2. For each parent Fab, three sub-libraries were constructed with different length and variability in the CDR H3 sequence: A11A, A11B and A11C for Fab A11 and E2A, E2B and E2C for Fab E2. The oligonucleotides used were A11-A, A11-Aa, A11-Ab, A11-B, A11-Ba, A11-Bb, A11-C, A11-Ca, A11-Cb, E2-A, E2-Aa, E2-Ab, E2-B, E2-Ba, E2-Bb, E2-C, E2-Ca, E2-Cb. A11A sub-library includes a direct graft of CDR H3 sequence from A11, with three different CDR H3 lengths, the wildtype length plus addition of one or two amino acids at the C-terminal end. In addition to the CDR H3 length variation, A11B sub-library keeps seven-residue turn GIAARRF constant, while A11C sub-library holds only RR motif constant. Aside from the one or two inserted codons, the codons encoding other residues in CDR H3 are soft randomized using a 70/10/10/10 nucleotide mix that on average allows for 50% of the wildtype amino acid and 50% of other amino acids. Three sub-libraries for E2 were constructed in the same way as that for A11.

The six mutagenesis reactions were electroporated separately into *Escherichia coli* SS320 and phage production was initiated by the addition of M13KO7 helper phage. After overnight growth at 37 °C, phage was harvested by precipitation with polyethylene glycol (PEG)/NaCl. The diversity of each sub-library and phage concentration after propagation are shown in **Supplement 1**. To check the quality of the libraries, mid-log phase *Escherichia coli* XL1-blue (Agilent Technologies) were infected with phage and the infected culture was plated on LB with 100 *µ*g/ml carbenicillin plate. Colonies from each sub-library were selected and the heavy chains were sequenced.

### 4.2 Selection against matriptase

The panning was done in three rounds as described previously. Each sub-library was screened separately (Schneider et al., 2012). Briefly, recombinant matriptase (Takeuchi et al., 1999) was adsorbed to the surface of a Nunc Maxisorp plate. Increasing stringency and decreasing protein concentration were conducted after each round, and individual colonies were selected, and Fab-phage were produced in a 96-well plate. Enzyme linked immunosorbent assay (ELISA) was performed with Fab-phage supernatant to verify binding of individual clones to matriptase. The sequences of the clones that bound matriptase were determined using VL Forward (5*^′^*-TGT AAA ACG ACG GCC AGT CTG TCA TAA AGT TGT CAC GG-3*^′^*) and VH Forward (5*^′^*-TGT AAA ACG ACG GCC AGT GGA CGC ATC GTG GCC CTA-3*^′^*) primers.

### 4.3 Biased Fab expression and purification

The phagemid DNA of ELISA positive clones were purified. A stop codon was introduced after the heavy chain constant domain using QuikChange^®^ Site-Directed Mutagenesis Kit (Stratagene). Fabs were then expressed in *Escherichia coli* 55244 cells (ATCC) as described previously (Paduch et al., 2013). Each Fab was freshly transformed to inoculate an overnight culture. The overnight culture was used to inoculate 1 L of low phosphate CRAP-Pi sAB expression media supplemented with 100 *µ*g/ml of ampicillin at 30 °C and 230 rpm. After 18 hours of growth, the bacteria were harvested and pelleted by centrifugation. The cells were resuspended in 40 ml of lysis buffer (B-PER from Pierce supplemented with 1 U/ml DNaseI). An equal volume of PBS pH 7.4 was added to the resuspended solution and the entire volume was transferred to a 250 ml centrifuge bottle. The solution was then heated at 65 °C for 30 min and centrifuged at 27,000*g* for 30 min. Fab supernatant was incubated with pre-washed Protein L Plus agarose (Pierce) for at least 1 h at 4 °C. Beads were spun down and purified over a gravity column. The column was washed with 100 mM phosphate pH 7.2, 0.15 M NaCl buffer, and the protein was eluted with 50 mM glycine pH 2 and immediately neutralized with 1 M Tris pH 9. Fabs were dialyzed against PBS pH 7.4 overnight, concentrated, and then purified with size exclusion chromatography. Fractions were collected and concentrated for storage at 4 °C.

### 4.4 Steady-state kinetics for matriptase inhibition

IC_50_ and relative *K*_I_ values were calculated for all the Fabs identified. The inhibition of matriptase and the kinetic parameters were calculated using the same protocols described before for A11 and E2 (Farady et al., 2007). Briefly, all reactions were carried out in 50 mM Tris pH 8.8, 50 mM NaCl, 0.01% Tween-20 in a 96-well, black, medium binding, flat-bottomed (Corning) where cleavage of substrate 200 *µ*M Spectrozyme^®^ tPA (hexahydrotyrosyl-Gly-Arg-pNA, American Diagnostica) was monitored in a UVMax Microplate Reader (Molecular Devices Corporation). IC_50_ values were determined by pre-incubating 0.2 nM matriptase with Fabs for 5 h to ensure steady-state kinetics. Relative *K*_I_ values were calculated from IC_50_ values according to *K*_I_ = IC_50_*/*(1 + [S]*/K*_m_) (Cheng and Prusoff, 1973).

All graphs and equations were fit using GraphPad Prism v7 (GraphPad Software).

### 4.5 Construction of TMPRSS2 expression vectors for biopanning

TMPRSS2 DNA was a generous gift from the Nelson lab (Lucas et al., 2014). Two extracellular constructs of TMPRSS2 were generated using this template and a potential glycosylation site was mutated to N249G. The first construct included the entire extracellular region (low-density lipoprotein LDL-receptor class A, scavenger receptor cysteine-rich SRCR, and serine protease) from amino acids 108–492; the second construct only contained the SRCR and serine protease domain from amino acids 150–492. A stop codon was inserted after the protease domain and no additional purification tags were included. These constructs were cloned into the pPICZ*α*B construct from Invitrogen’s EasySelect *Pichia* Expression Kit according to the manufacturer’s protocol, transformed into *Pichia pastoris* strain X33, and selected on plates containing increasing concentrations of Zeocin to identify multi-copy recombinants. Clones were screened via small scale expression measuring TMPRSS2 activity in the supernatant daily with a chromogenic substrate after induction.

### 4.6 TMPRSS2 expression and purification

The expression and purification were modified from a previous report (Lucas et al., 2014). Briefly, a single colony was grown in 10 ml buffered medium with glycerol (BMGY) overnight at 30 °C and 230 rpm. The overnight culture was used to inoculate 1 L BMGY and grown until OD 2–6. Cells were pelleted and resuspended in 100 ml buffered medium with methanol (BMMY). Cells were induced with 5% methanol every 24 h for 72–96 h. To harvest TMPRSS2 protein, cells were centrifuged and the supernatant was precipitated with 70% ammonium sulfate at 4 °C overnight. Protein was harvested at 27,000 × *g* for 45 min, resuspended in 50 mM Tris pH 8, 0.5 M NaCl, 0.01% CHAPS, and solubilized for 2 h at 4 °C. The solubilized protein was then purified over a gravity column containing soybean trypsin inhibitor immobilized agarose (Pierce). The column was washed with 50 mM Tris pH 8, 150 mM NaCl buffer and the protein was eluted with 50 mM glycine pH 2 and neutralized 1:10 with 1 M Na-acetate. TMPRSS2 was concentrated and then purified with size exclusion chromatography using 50 mM potassium phosphate buffer pH 6, 150 mM NaCl, 1% glycerol to prevent auto-proteolysis. Fractions were collected, concentrated and stored at 4 °C for immediate use or flash frozen with liquid nitrogen and stored at −80 °C for longer term.

### 4.7 Identification of Fabs against TMPRSS2

Recombinant TMPRSS2 was biochemically biotinylated with EZ-Link^®^ NHS-Chromogenic Biotinylation Kit (Thermo Scientific) to obtain 1–2 biotin molecules per protein and then buffer exchanged over a PD-10 desalting column. The automated selection with Library F was conducted on streptavidin beads as described previously (Hornsby et al., 2015). Binding selections with the biased and naïve libraries were conducted with active biotinylated TM-PRSS2 immobilized to streptavidin-coated wells in Nunc Maxisorp plates. Three rounds of selection with decreasing protein concentration (20, 10, and 5 *µ*g/ml) and increasing wash stringency were completed. After enrichment in the third round, single colonies were selected, and Fab-phage supernatant or Fab supernatant for biased and naïve library respectively was used in ELISA to identify TMPRSS2 positive clones with 1:5000 anti-M13 peroxidase (GE Healthcare Life Sciences) and 1:1000 anti-c-myc peroxidase (Roche) used as a secondary antibody. Semi-quantitative ELISA was performed using serial dilutions of the supernatant to prioritize individual clones as described previously (Kim et al., 2011). Individual clones were sequenced with VL Forward and VH Forward primers for the biased library and JS-1 (5*^′^*-AGC GGA TAA CAA TTT CAC ACA GG-3*^′^*) and JS-2 (5*^′^*-TTT GTC GTC TTT CCA GAC GTT AGT-3*^′^*) primers for the naïve library.

### 4.8 Expression and purification of Fabs from naïve library

Fabs from the naïve library were expressed in *Escherichia coli* BL21-Gold (DES) and purified as previously described (Kim et al., 2011). A single colony from a fresh plate was used to inoculate an overnight culture. The overnight culture was used to inoculate 1 L of 2×YT with 0.1% glucose and 100 *µ*g/ml ampicillin and grown until log phase. When the OD_600_ reached 0.5–0.8, the cells were induced with 1 mM IPTG and grown at 20 °C at 200 rpm overnight. The bacteria were harvested and pelleted by centrifugation. The pellet was subjected to osmotic shock for periplasmic protein purification. The periplasmic fraction was loaded onto a gravity column with Ni-NTA resin (Thermo Scientific) and washed. Protein was eluted with buffer containing 500 mM imidazole. Protein was dialyzed against PBS pH 7.4 overnight at 4 °C. Fabs were concentrated and then purified with size exclusion chromatography. Fractions were collected and concentrated for storage at 4 °C.

### 4.9 Second generation biased library construction and affinity maturation selections

The library was constructed as described previously (Chen and Sidhu, 2014). Phage pools from a small set of biased hits (biAb-TM-1, biAb-TM-7, biAb-TM-8, and biAb-TM-5) were used to infect CJ236 cells to create dU-ssDNA. A set of mutagenic oligonucleotides were designed to introduce diversity at CDR L1 and CDR H1: L1-11, L1-12, L1-16, L1-17, H1-7, H1-8, H1-9, ordered from Trilink Biotechnologies (San Diego, CA). CCC-dsDNA was transformed into electrocompetent SS320 co-infected with helper phage M13K07 (NEB). To check the mutation rate of the library, mid-log phase *Escherichia coli* TG1 cells (Lucigen) were infected with phage from the library and plated on LB with 100 *µ*g/ml ampicillin plate. Colonies were selected and the heavy and light chains were sequenced.

Two additional rounds of selection against TMPRSS2 with the affinity maturation library were conducted. In each round, decreasing protein concentration (1 *µ*g/ml, and 0.1–0.5 *µ*g/ml) and increasing wash stringency were used to select for more potent Fabs. Individual clones were characterized as described in the previous section.

### 4.10 *K*_D_ determination for Fabs

Kinetic constants for each Fab were determined using an Octet RED384 instrument (ForteBio). Three concentrations of each Fab (10 *µ*M, 1 *µ*M, and 100 nM) were tested for binding to biotinylated TMPRSS2 immobilized on a Streptavidin SA biosensor (ForteBio). All measurements were performed at room temperature in black Grenier 384-well microplates in running buffer PBS pH 7.4 with 0.1% (w/v) bovine serum albumin (BSA) and 0.02% (v/v) Tween-20. Biotinylated TMPRSS2 was loaded for 100 s from a 50 nM solution onto the biosensor and then the baseline was equilibrated for 60 s in buffer. The association phase for Fabs was 300 s followed by 120 s dissociation in buffer. Between each Fab sample, the biosensors were regenerated in three cycles of 10 mM glycine pH 1.5 for 5 s followed by buffer for 5 s. Data were analyzed using a 1:1 interaction model on the ForteBio data analysis software 8.2.

### 4.11 biAb-TM-5 mutant construction and inhibition assays

Point mutants to alanine and other residues were made from the wildtype plasmid using QuikChange^®^ Site-Directed Mutagenesis Kit (NEB) and oligos purchased from Azenta. These mutants were expressed as described previously and their inhibitory potency was measured against 30 nM commercial TMPRSS2 (CusaBio) at a standardized 5 *µ*M Fab concentration with Boc-QAR-AMC substrate (R&D Systems) at 250 *µ*M. Reactions were carried out in 30 *µ*L of 50 mM Tris pH 8, 50 mM NaCl, 0.01% Tween-20 and reaction rates measured with a BioTek H4 plate reader.

### 4.12 Fab mutant ΔΔ*G* energy estimation

Using Fab–protease complex models generated from AlphaFold2, the ΔΔ*G* binding energy was predicted for all possible residue substitutions across each of the CDR loop contacts. This was conducted for biased Fab biAb-TM-5 in complex with TMPRSS2 using the modeled complex structure from AlphaFold 2 multimer. No other modifications were made to the AF2 models besides the standard AMBER minimization. The structure used for Rosetta Flex ΔΔ*G* was additionally prepared using the standard FastRelax protocol for efficient use within Rosetta.

ΔΔ*G* scores from FoldX were generated for all antibody interface residues using the PSSM program which bundles BuildModel and AnalyseComplex methods in FoldX 5.0 (Delgado et al., 2019). Scores from Rosetta were obtained from the Flex ΔΔ*G* application with default flags (nstruct = 35) from the authors using Rosetta 3.12 (Barlow et al., 2018). mCSM-AB2 (Myung et al., 2020) predictions were conducted using the webserver (https://biosig.lab. uq.edu.au/mcsm_ab2/prediction); standard options were used, and saturation mutagenesis was selected. GeoPPI (Liu et al., 2021b) and GearBind (Cai et al., 2024) were run for saturating mutagenesis of all interface residues with default options for each method.

Scores from each ΔΔ*G* prediction method were first sign-adjusted to match a common stability convention (negative = stabilizing) and then normalized using global maximum absolute scaling. The resulting values were subsequently averaged across methods to generate a consensus ΔΔ*G* profile heatmap for saturating mutations at each interface CDR position.

### 4.13 Modeling of Fab–protease complexes

AlphaFold2 models used for ΔΔ*G* predictions were generated with version 2.3.2 (multi-mer v3) (Evans et al., 2022; Jumper et al., 2021) from (https://github.com/deepmind/alphafold) and run on the UCSF Wynton shared computing environment. Raw MSAs were prepared for each sequence input using the full AlphaFold pipeline and querying the full databases (UniRef90 2020 01, MGnify 2018 12, Uniclust 2018 8 and BFD). Default options with amber relaxation were used. Templates were disabled for all predictions to assess genuine predictive capability rather than template-based modeling; separate runs with templates enabled are reported in **Supplement 3**. AF2 generates 5 models for each input which are ranked by pTM score; unless otherwise noted, the top-ranked model was used for analysis.

ColabFold (AlphaFold2-multimer) models for benchmarking comparison between A11, E2, and AB2 were generated using LocalColabFold 1.5.5 (Mirdita et al., 2022) with AMBER relax and 24 recycles. For each target, 5 seeds were run with 5 models per seed, yielding 25 predictions per complex. Templates were disabled for primary analysis; runs with templates enabled are reported in **Supplement 3**. The top-ranked model by pTM score was used for analysis.

AlphaFold3 models were generated using a local installation of AlphaFold3 (Abramson et al., 2024). For each target, 100 seeds were run with 5 samples per seed, yielding 500 predictions per complex. Templates were disabled for primary analysis; runs with templates enabled are reported in **Supplement 3**. The top-ranked model by ipTM score was used for analysis.

Boltz-2 models were generated using Boltz-2 v2.2.1 (https://github.com/jwohlwend/boltz) (Passaro et al., 2025). For each target, 100 seeds were run yielding one model per seed (100 predictions per complex). The top-ranked model by Boltz-2 confidence score was used for analysis.

Chai-1 models were generated using Chai-1 v0.6.1 (https://github.com/chaidiscovery/chai-lab) (Discovery et al., 2024). For each target, 100 seeds were run with 5 models per seed, yielding 500 predictions per complex. The top-ranked model by Chai-1 confidence score was used for analysis.

OpenFold3 models were generated using OpenFold3-preview v0.3.1 (https://github. com/aqlaboratory/openfold-3) (The OpenFold3 Team, 2025). For each target, 5 seeds were run with 5 samples per seed, yielding 25 predictions per complex.

Higher seed counts (100) were used for AF3, Boltz-2, and Chai-1 to assess confidence calibration and sampling variance across prediction replicates from the latest diffusion-based modeling methods; seed convergence analysis is reported in **Supplement 3, continued**.

### 4.14 ClusPro rigid-body docking

ClusPro 2.0 (Kozakov et al., 2017; Vajda et al., 2017) was used for rigid-body docking of experimentally determined structures. For each target complex (A11–matriptase PDB: 3SO3, E2–matriptase PDB: 3BN9, and AB2–uPA PDB: 9PYF), the Fab and protease domain chains were extracted from the deposited crystal structures and submitted as separate receptor and ligand inputs to the ClusPro web server (https://cluspro.org). The antibody mode was selected. ClusPro generated 25–30 cluster representatives per target. ClusPro docking used PDB-deposited structures as rigid-body inputs, whereas co-folding methods used input sequences only.

### 4.15 DockQ scoring

All complex models were scored against reference crystal structures using DockQ v2 (Mirabello and Wallner, 2024). DockQ scores reflect an average of the fraction of native contacts (*f*_nat_), scaled interface RMSD (iRMSD), and scaled ligand RMSD (LRMSD). Models were classified according to CAPRI quality thresholds: Incorrect (DockQ *<* 0.23), Acceptable (0.23–0.49), Medium (0.49–0.80), and High Quality (*>* 0.80). Reference structures used were A11–matriptase (PDB: 3SO3), E2–matriptase (PDB: 3BN9), and AB2–uPA (PDB: 9PYF).

All structures shown in figures are rendered in ChimeraX (UCSF).

## Ethical and legal declarations

## Author contributions

Kyle J. Anderson: conceptualization – computational portion, writing – evolved manuscript, formal analysis, data curation, validation, visualization, software, investigation, methodology; Melody S. Lee: conceptualization – biased libraries, writing – initial draft, investigation, methodology; Natalia Sevillano: investigation; Gang Chen: methodology, supervision, resources; Michael J. Hornsby: methodology, supervision, resources; Sachdev S. Sidhu: resources, supervision, methodology; Charles S. Craik: supervision, project administration, funding acquisition, resources, review and editing.

## Research funding

NIH training grant [5T32GM008284-33 to K.J.A.]; HARC center: National Institutes of Health [U54AI170792 to C.S.C.]; National Institute of Allergy and Infectious Disease AViDD [U19AI171110 to C.S.C.].

## Abbreviations

AF2: AlphaFold2
AF3: AlphaFold3
BLI: biolayer interferometry
CDR: complementarity determining region
CAPRI: Critical Assessment of Predicted Interactions
Fab: fragment antigen binding
H3: heavy chain CDR loop 3
IC_50_: half-maximal inhibitory concentration
*K*_D_: dissociation constant
*K*_I_: inhibition constant
L1: light chain CDR loop 1
MT-SP1: membrane-type serine protease 1 (matriptase)
PAE: predicted aligned error
pLDDT: predicted local distance difference test
TTSP: type II transmembrane serine pro-tease
TMPRSS2: transmembrane protease serine 2
uPA: urokinase plasminogen activator

## Supplementary figure legends

**Supplement 1: Biased Library Construction.** (A) Structure of A11 in complex with matriptase (PDB: 3SO3) with CDRs (L3, H1, H2, H3) annotated along with the degenerate codon oligos used to generate six biased sub-libraries from parental Fabs A11 and E2. (B) CDR sequences for biased library hits against matriptase with corresponding *K*_I_ values; residues are colored by amino acid properties to highlight diversity across CDR L3, H1, H2, and H3. (C) Library diversity and phage concentration (cfu/mL) for each sub-library.

**Supplement 2: Affinity Maturation.** Affinity maturation of biAb-TM-5: (A) AlphaFold2 structure of Fab 41D1 (light chain, pink; heavy chain, green) with CDR L1 and H1 annotated, shown aligned with crystal structures of matriptase (PDB: 3SO3, orange) and TM-PRSS2 (PDB: 7MEQ, blue), alongside an AlphaFold2-predicted TMPRSS2 model. CDR L1 and H1 diversification oligos used for affinity maturation are shown in the accompanying tables. (B) Inhibition curves for anti-TMPRSS2 clones biAb-TM-7 (642 nM), biAb-TM-5 (IC_50_ 2.8 *µ*M), biAb-TM-8 (IC_50_ 40 *µ*M), and affinity-matured clones biAb-TM-5.1 (472 nM) and biAb-TM-5.2 (376 nM); TMPRSS2 full extracellular domain at 17.5 pM final concentration with Boc-QAR-AMC substrate. (C) BLI binding curves for affinity-matured TMPRSS2 Fabs (clones 1C1, 1B5, 1F12, 1F5, 5G6, 5H1) at concentrations ranging from 10 *µ*M to 1 *µ*M.

**Supplement 3: Detailed AlphaFold Metrics.** (A) DockQ scores comparing rank-1 (top-ranking) versus top-scoring models across co-folding methods (ColabFold and AF3) with and without templates, evaluated against reference structures for each antibody–protease complex. (B) Structure prediction versus docking method comparison, showing DockQ scores from co-folding methods (Chai-1, ColabFold, AF3) alongside traditional docking approaches (ClusPro, HDOCK, HADDOCK) for each target complex. (C) pLDDT confidence metric and predicted aligned error (PAE) from the AF3 model of PDB 3SO3 (A11–matriptase), highlighting CDR H3 loop conformational differences between the predicted model and the crystal structure.

**Supplement 3 (continued): Co-folding Benchmark and Seed Convergence.** (A) Best (per-seed best) versus confidence-selected DockQ scores for five co-folding methods (AF3, Boltz-2, Chai-1, OpenFold3, ColabFold) across four antibody–protease complexes: A11–matriptase (PDB: 3SO3), E2–matriptase (PDB: 3BN9), AB2–uPA using the exact PDB-deposited sequence (PDB: 9PYF), and AB2–uPA scored against an active conformation reference. Annotations indicate the difference between top-scoring and confidence-selected DockQ; the dashed green line marks the 0.80 DockQ threshold for high-quality models. (B) Seed convergence analysis showing cumulative best DockQ score as a function of the number of random seeds sampled for AF3, Boltz-2, and Chai-1, with shaded regions indicating variability across bootstrap replicates.

**Supplement 4: Per-method** ΔΔ*G* **Reports.** (A) Scaled ΔΔ*G* heatmaps for each of the five computational methods (FoldX, mCSM-AB2, Rosetta, GeoPPI, GearBind), showing predicted stability changes for all single-point mutations across antibody CDR positions. Red indicates destabilizing mutations and blue indicates stabilizing mutations. (B) ΔΔ*G* data coverage for each antibody position across each method, with darker shading indicating more complete mutational coverage. (C) Number of antibody positions covered by each method, summarizing how many of the five methods provide predictions at each position.

**Supplement 5: Detailed** ΔΔ*G* **Reports.** (A) Correlation matrix for ΔΔ*G* values between the five methods (FoldX, mCSM-AB2, Rosetta, GeoPPI, GearBind) computed on common mutations, highlighting agreement and divergence between predictors. (B) Distribution of scaled ΔΔ*G* values for each method shown as violin plots. (C) Distribution of all averaged (consensus) ΔΔ*G* values across mutations, with the neutral (ΔΔ*G* = 0) threshold marked. (D) Average ΔΔ*G* across antibody CDR positions, identifying positions with consistently favorable or unfavorable predicted stability changes. (E) Per-method mean ΔΔ*G* heatmap for each antibody position (Chothia numbering), enabling direct comparison of method-level predictions at each residue.

**Supplement 6:** ΔΔ*G* **vs. Biopanning Data.** (A) Heatmaps showing amino acid frequencies recovered from biopanning selection, ΔΔ*G* predictions by position and mutation, and the overlay of ΔΔ*G* against biopanning frequency. (B) Correlation between consensus ΔΔ*G* values and biopanning residue frequencies (Pearson *r*), assessing the predictive relationship between computational stability estimates and experimental selection outcomes. (C) ΔΔ*G* range by position across mutated CDR residues. (D) Comparison of best observed ΔΔ*G* for mutations to residues with the same chemistry versus amino acid conservation frequency from biopanning, with residues categorized as highly favorable, slightly enhancing, neutral, or ≥1*σ* pruned. (E) ΔΔ*G* distributions for residues observed versus not observed in biopanning selection. (F) ΔΔ*G* distribution stratified by biopanning frequency category (low, medium-low, medium-high, and high frequency). (G) Enrichment analysis by ΔΔ*G* category, comparing observed versus expected residue counts from biopanning. (H) ΔΔ*G* distributions stratified by chemistry change type (conservative, moderate, radical). (I) ΔΔ*G* distributions with chemistry-adjusted groupings, accounting for the physicochemical similarity of substitutions.

