## Supplementary Material for "Structural Basis of Serine Protease Inhibition by Antibodies from Biased Fab Phage-Display Libraries"

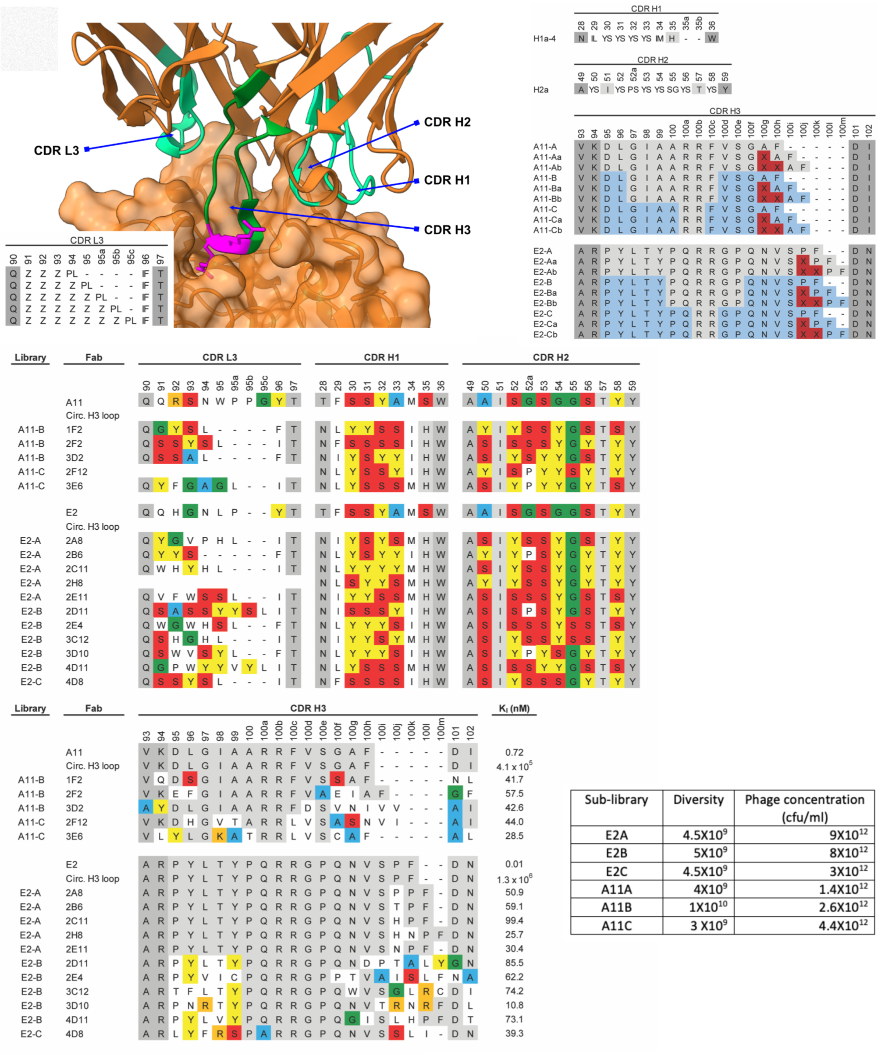

| Sub-library | Diversity | Phage concentration (cfu/ml) |
|---|---|---|
| E2A | $4.5X10^9$ | $9X10^{12}$ |
| E2B | $5X10^9$ | $8X10^{12}$ |
| E2C | $4.5X10^9$ | $3X10^{12}$ |
| A11A | $4X10^9$ | $1.4X10^{12}$ |
| A11B | $1X10^{10}$ | $2.6X10^{12}$ |
| A11C | $3 X10^9$ | $4.4X10^{12}$ |

X indicates trimer mix
Y indicates C, T wobble site
W indicates A, T wobble site
M indicates A, C wobble site
R indicates A, G wobble site
S indicates C, G wobble site
Lower case = soft randomization with 70/10/10/10 nucleotide mix
VNT codon ([GAC][GACT]T)
ZZZ indicates amino acid mixture with 11:10:4:3:3:2:2:2:1:1:1 (S/Y/G/W/D/A/H/P/F/V/R)

Supplemental Figure 1: **Biased Library Construction.** (A) Structure of A11 in complex with matriptase (PDB: 3SO3) with CDRs (L3, H1, H2, H3) annotated along with the degenerate codon oligos used to generate six biased sub-libraries from parental Fabs A11 and E2. (B) CDR sequences for biased library hits against matriptase with corresponding $K_I$ values; residues are colored by amino acid properties to highlight diversity across CDR L3, H1, H2, and H3. (C) Library diversity and phage concentration (cfu/mL) for each sub-library.

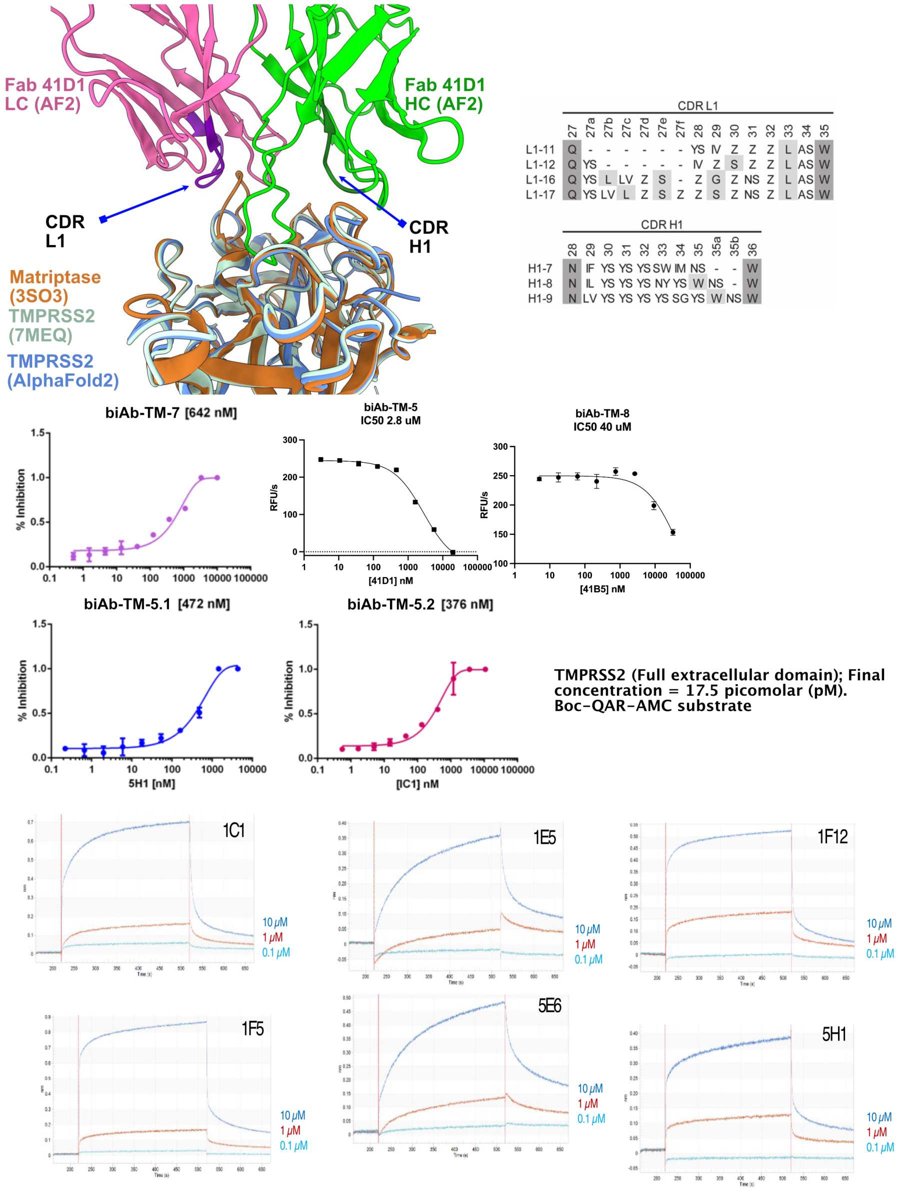

| CDR L1 | 27 | 27a | 27b | 27c | 27d | 27e | 27f | 28 | 29 | 30 | 31 | 32 | 33 | 34 | 35 |
|---|---|---|---|---|---|---|---|---|---|---|---|---|---|---|---|
| L1-11 | Q | - | - | - | - | - | - | YS | IV | Z | Z | Z | L | AS | W |
| L1-12 | Q | YS | - | - | - | - | - | IV | Z | S | Z | Z | L | AS | W |
| L1-16 | Q | YS | L | LV | Z | S | - | Z | G | Z | NS | Z | L | AS | W |
| L1-17 | Q | YS | LV | L | Z | S | Z | Z | S | Z | NS | Z | L | AS | W |

| CDR H1 | 28 | 29 | 30 | 31 | 32 | 33 | 34 | 35 | 35a | 35b | 36 |
|---|---|---|---|---|---|---|---|---|---|---|---|
| H1-7 | N | IF | YS | YS | YS | SW | IM | NS | - | - | W |
| H1-8 | N | IL | YS | YS | YS | NY | YS | W | NS | - | W |
| H1-9 | N | LV | YS | YS | YS | YS | SG | YS | W | NS | W |



Supplemental Figure 2: **Affinity Maturation of biAb-TM-5.** (A) AlphaFold2 structure of Fab 41D1 (light chain, pink; heavy chain, green) with CDR L1 and H1 annotated, shown aligned with crystal structures of matriptase (PDB: 3SO3, orange) and TMPRSS2 (PDB: 7MEQ, blue), alongside an AlphaFold2-predicted TMPRSS2 model. CDR L1 and H1 diversification oligos used for affinity maturation are shown in the accompanying tables. (B) Inhibition curves for anti-TMPRSS2 clones biAb-TM-7 (642 nM), biAb-TM-5 ($IC_{50}$ 2.8 $\mu$M), biAb-TM-8 ($IC_{50}$ 40 $\mu$M), and affinity-matured clones biAb-TM-5.1 (472 nM) and biAb-TM-5.2 (376 nM); TMPRSS2 full extracellular domain at 17.5 pM final concentration with Boc-QAR-AMC substrate. (C) BLI binding curves for affinity-matured TMPRSS2 Fabs (clones 1C1, 1B5, 1F12, 1F5, 5G6, 5H1) at concentrations ranging from 10 $\mu$M to 1 $\mu$M.

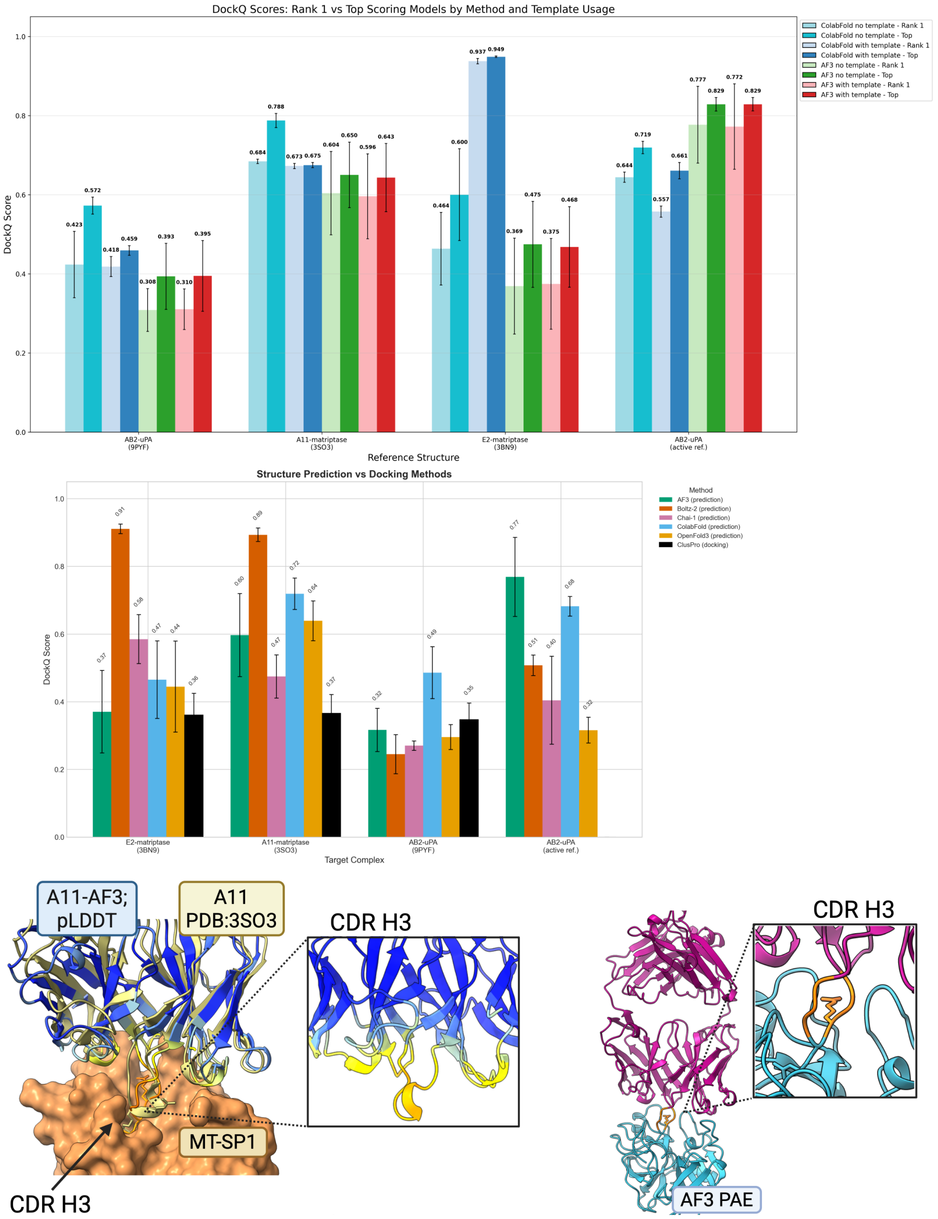


Supplemental Figure 3: **Detailed AlphaFold Metrics.** (A) DockQ scores comparing rank-1 (top-ranking) versus top-scoring models across co-folding methods (ColabFold and AF3) with and without templates, evaluated against reference structures for each antibody–protease complex. (B) Structure prediction versus docking method comparison, showing DockQ scores from co-folding methods (Chai-1, ColabFold, AF3) alongside traditional docking approaches (ClusPro, HDOCK, HADDOCK) for each target complex. (C) pLDDT confidence metric and predicted aligned error (PAE) from the AF3 model of PDB 3SO3 (A11–matriptase), highlighting CDR H3 loop conformational differences between the predicted model and the crystal structure.

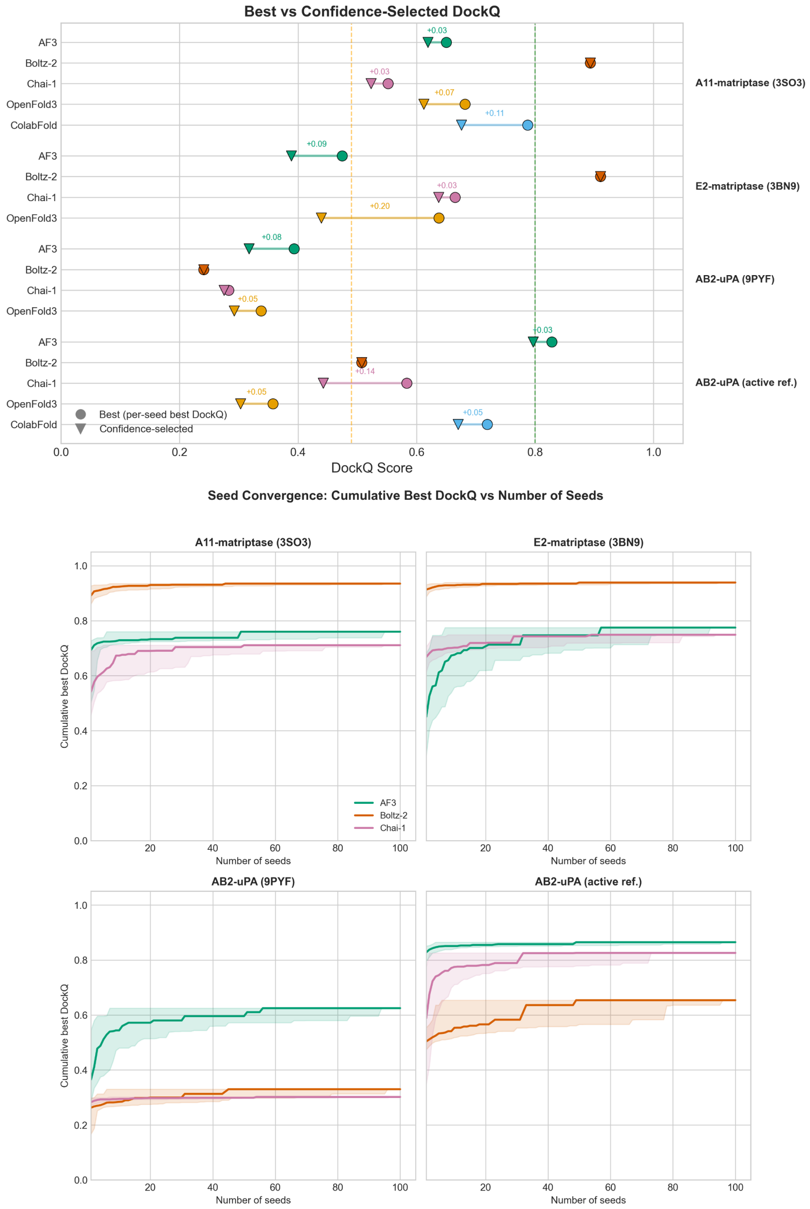


Supplemental Figure 4: **Detailed AlphaFold Metrics (continued): Co-folding Benchmark and Seed Convergence.** (A) Best (per-seed best) versus confidence-selected DockQ scores for five co-folding methods (AF3, Boltz-2, Chai-1, OpenFold3, ColabFold) across four antibody–protease complexes: A11–matriptase (PDB: 3SO3), E2–matriptase (PDB: 3BN9), AB2–uPA using the exact PDB-deposited sequence (PDB: 9PYF), and AB2–uPA scored against an active conformation reference. Annotations indicate the difference between top-scoring and confidence-selected DockQ; the dashed green line marks the 0.80 DockQ threshold for high-quality models. (B) Seed convergence analysis showing cumulative best DockQ score as a function of the number of random seeds sampled for AF3, Boltz-2, and Chai-1, with shaded regions indicating variability across bootstrap replicates. Results demonstrate how rapidly each method converges to its best achievable DockQ for each complex.

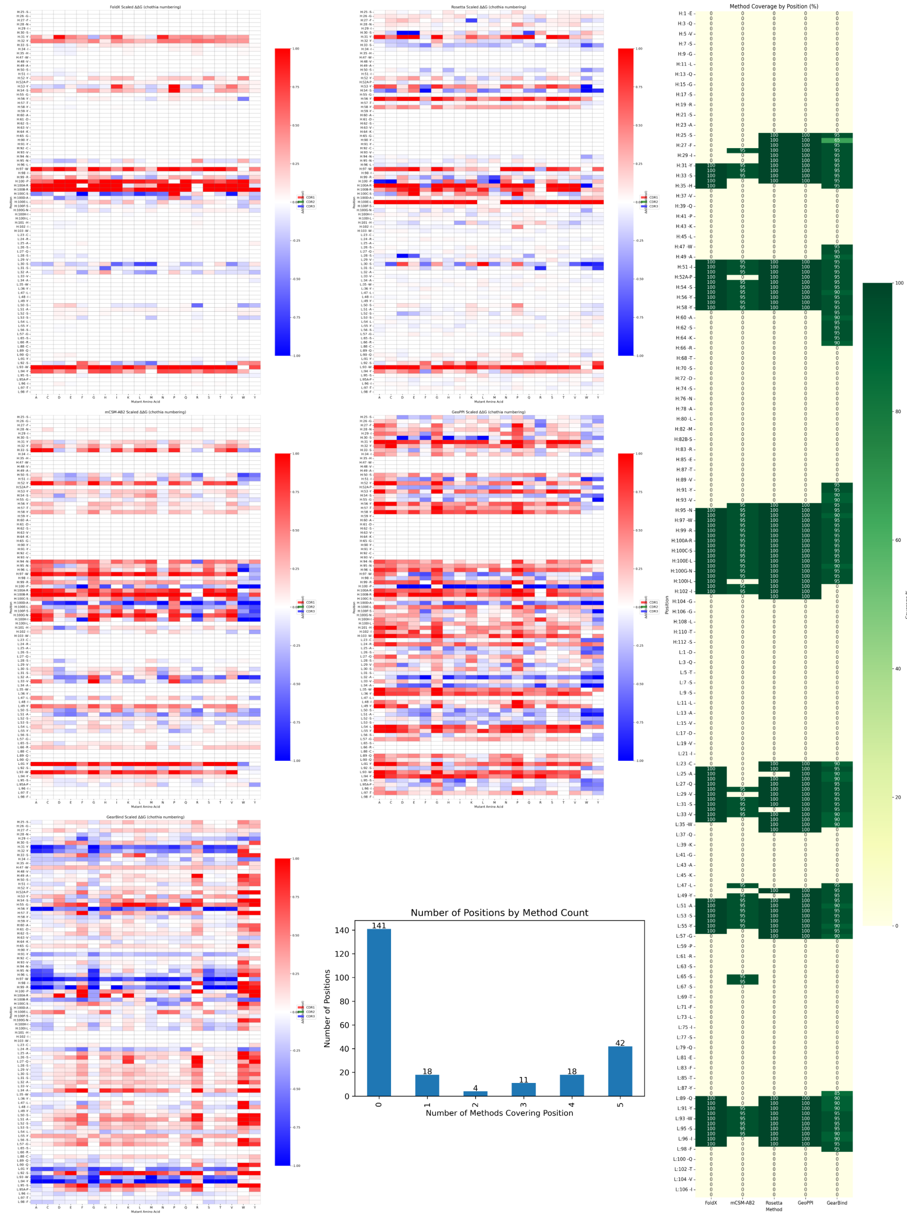


Supplemental Figure 5: **Per-method $\Delta\Delta G$ Reports.** (A) Scaled $\Delta\Delta G$ heatmaps for each of the five computational methods (FoldX, mCSM-AB2, Rosetta, GeoPPI, GearBind), showing predicted stability changes for all single-point mutations across antibody CDR positions. Red indicates destabilizing mutations and blue indicates stabilizing mutations. (B) $\Delta\Delta G$ data coverage for each antibody position across each method, with darker shading indicating more complete mutational coverage. (C) Number of antibody positions covered by each method, summarizing how many of the five methods provide predictions at each position.

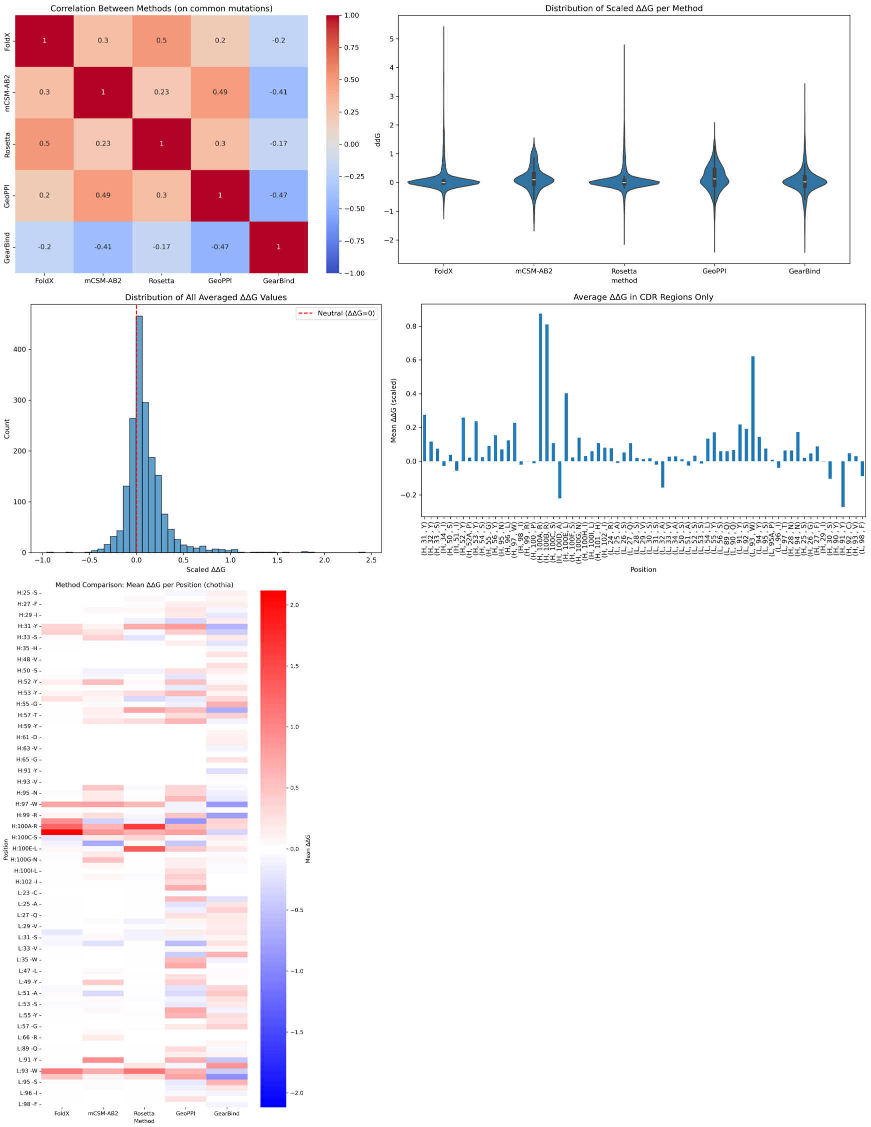


Supplemental Figure 6: **Detailed $\Delta\Delta G$ Reports.** (A) Correlation matrix for $\Delta\Delta G$ values between the five methods (FoldX, mCSM-AB2, Rosetta, GeoPPI, GearBind) computed on common mutations, highlighting agreement and divergence between predictors. (B) Distribution of scaled $\Delta\Delta G$ values for each method shown as violin plots. (C) Distribution of all averaged (consensus) $\Delta\Delta G$ values across mutations, with the neutral ($\Delta\Delta G = 0$) threshold marked. (D) Average $\Delta\Delta G$ across antibody CDR positions, identifying positions with consistently favorable or unfavorable predicted stability changes. (E) Per-method mean $\Delta\Delta G$ heatmap for each antibody position (Chothia numbering), enabling direct comparison of method-level predictions at each residue.

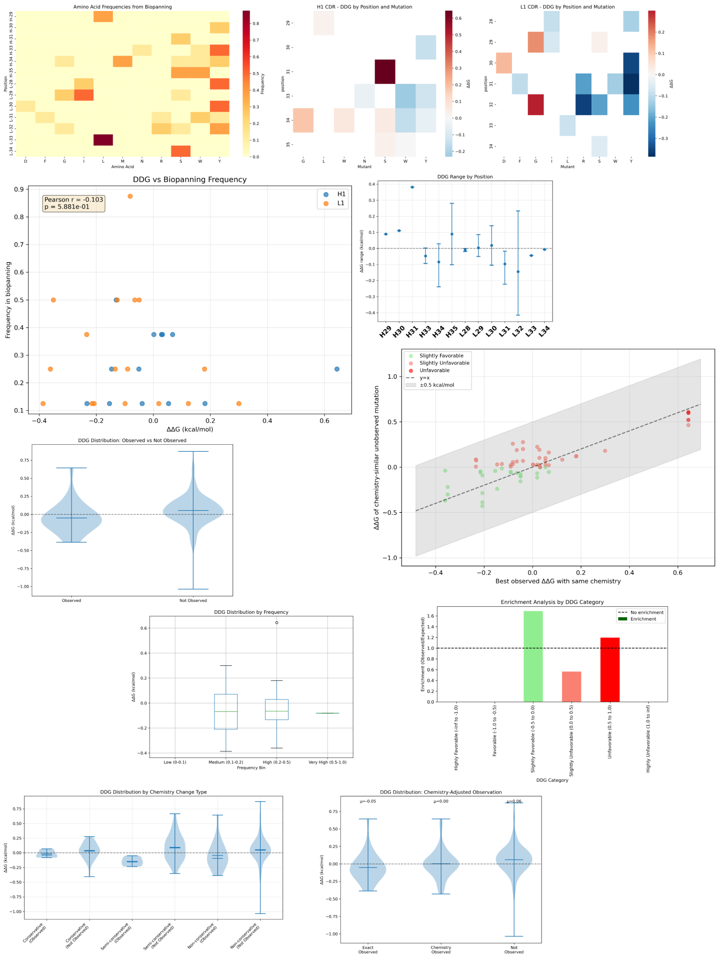


Supplemental Figure 7: $\Delta\Delta G$ **vs. Biopanning Data.** (A) Heatmaps showing amino acid frequencies recovered from biopanning selection, $\Delta\Delta G$ predictions by position and mutation, and the overlay of $\Delta\Delta G$ against biopanning frequency. (B) Correlation between consensus $\Delta\Delta G$ values and biopanning residue frequencies (Pearson $r$), assessing the predictive relationship between computational stability estimates and experimental selection outcomes. (C) $\Delta\Delta G$ range by position across mutated CDR residues. (D) Comparison of best observed $\Delta\Delta G$ for mutations to residues with the same chemistry versus amino acid conservation frequency from biopanning, with residues categorized as highly favorable, slightly enhancing, neutral, or $\geq 1\sigma$ pruned. (E) $\Delta\Delta G$ distributions for residues observed versus not observed in biopanning selection. (F) $\Delta\Delta G$ distribution stratified by biopanning frequency category (low, medium-low, medium-high, and high frequency). (G) Enrichment analysis by $\Delta\Delta G$ category, comparing observed versus expected residue counts from biopanning. (H) $\Delta\Delta G$ distributions stratified by chemistry change type (conservative, moderate, radical). (I) $\Delta\Delta G$ distributions with chemistry-adjusted groupings, accounting for the physicochemical similarity of substitutions.
